# Discovery of Aminoglycosides as first in class, nanomolar inhibitors of Heptosyltransferase I

**DOI:** 10.1101/2021.10.03.462930

**Authors:** Jozafina Milicaj, Bakar Hassan, Joy M. Cote, Carlos A. Ramirez-Mondragon, Nadiya Jaunbocus, Angelika Rafalowski, Kaelan R. Patel, Colleen D. Castro, Ramaiah Muthyala, Yuk Y. Sham, Erika A. Taylor

## Abstract

A clinically relevant inhibitor for Heptosyltransferase I (HepI) has been sought after for many years and while many have discovered or designed novel small molecule inhibitors, these compounds lack the bioavailability and potency necessary for therapeutic use. Extensive characterization of the HepI protein has provided valuable insight into the dynamic motions necessary for catalysis that could be targeted for inhibition. Structural similarity inspection of Kdo_2_-lipid A suggested aminoglycoside antibiotics as potential inhibitors for HepI. Multiple aminoglycosides have been experimentally validated to be first-in-class nanomolar inhibitors, with the best inhibitor demonstrating a *K*_*i*_ of 600 +/− 90 nM. Detailed kinetic analyses were performed to determine the mechanism of inhibition while circular dichroism spectroscopy, intrinsic tryptophan fluorescence, docking, and molecular dynamics simulations were used to corroborate kinetic experimental findings. Kinetic analysis methods include Lineweaver-Burk, Dixon, Cornish-Bowden and Mixed-Model of Inhibition which allowed for unambiguous assignment of inhibition mechanism for each inhibitor. In this study, we show that neomycin and kanamycin b are competitive inhibitors against the sugar acceptor substrate while tobramycin exhibits a mixed inhibitory effect and streptomycin is non-competitive. Molecular dynamics simulations also allowed us to suggest that the inhibitors bind tightly and inhibit native dynamics due to a major desolvation penalty of the enzyme active site. While aminoglycosides have long been known as a class of potent antibiotics targeting bacterial ribosomes’ protein synthesis, they also have been scientifically shown to impact cell membrane stability. Our discovery suggest an additional novel mechanism of action of aminoglycosides in the inhibition of HepI which may provide a viable repurposing strategy of an available drug for disrupting lipopolysaccharide biosynthesis.

## Introduction

The ever-growing number of infections and deaths due to antibiotic resistant bacteria is a global health issue necessitating development of novel therapeutics to treat these new multi-drug resistant species. Gram-negative bacteria are more likely to develop resistance than Gram-positive bacteria (1), in part due to their complex membrane morphology and the extracellular polymeric substances (EPS) which decrease the permeability of xenobiotics and improve surface adhesion (2). EPS provide structural integrity to the intracellular matrix of a bacterial biofilm, which further enhances their resistance to hydrophobic antibiotics (3) (4). Lipopolysaccharide (LPS) is a major component of the EPS making up approximately 30% of the outer membrane leaflet while facilitating multiple purposes including enabling cellular motility, adhesion, and nutrient retrieval (5) (6). Cellular exposure to an extracellular threat such as an antibiotic or antigen, can trigger modification of LPS to further fortify the outer membrane leaflet, a common resistance mechanism known to reduce membrane permeability (1) (7) (8).

Previous studies have shown that when the LPS pathway genes are knocked out, it causes increased susceptibility to antibiotics, reduced cellular motility, reduced cellular adhesion, and in some cases was fatal to the cell (9). This information taken with the knowledge that the inner oligosaccharide core biosynthesis of LPS is conserved among all Gram-negative bacteria, reveals a potential target that can be exploited for therapeutic design (10). The first step in the LPS core biosynthetic pathway is catalyzed by Heptosyltransferase I (HepI), also known as WaaC (or RfaC), which is responsible for the transfer of a heptose moiety onto the first Kdo of Kdo_2_-Lipid A (Kdo: 3-deoxy-D-*manno*-oct-2-ulosonic acid) (11) (Figure 1a). HepI is a GT-B glycosyltransferase enzyme with two Rossman-like domains (connected by a linker region), which each bind one of the two enzymatic substrates (the N-terminal domain binds the nucleophilic Kdo_2_-Lipid A, while the C-terminal domain binds the electrophilic sugar donor ADP-L-*glycero*-β-D-*manno*-heptose, or ADP-Hep) (Figure1b).

**Figure 1:**
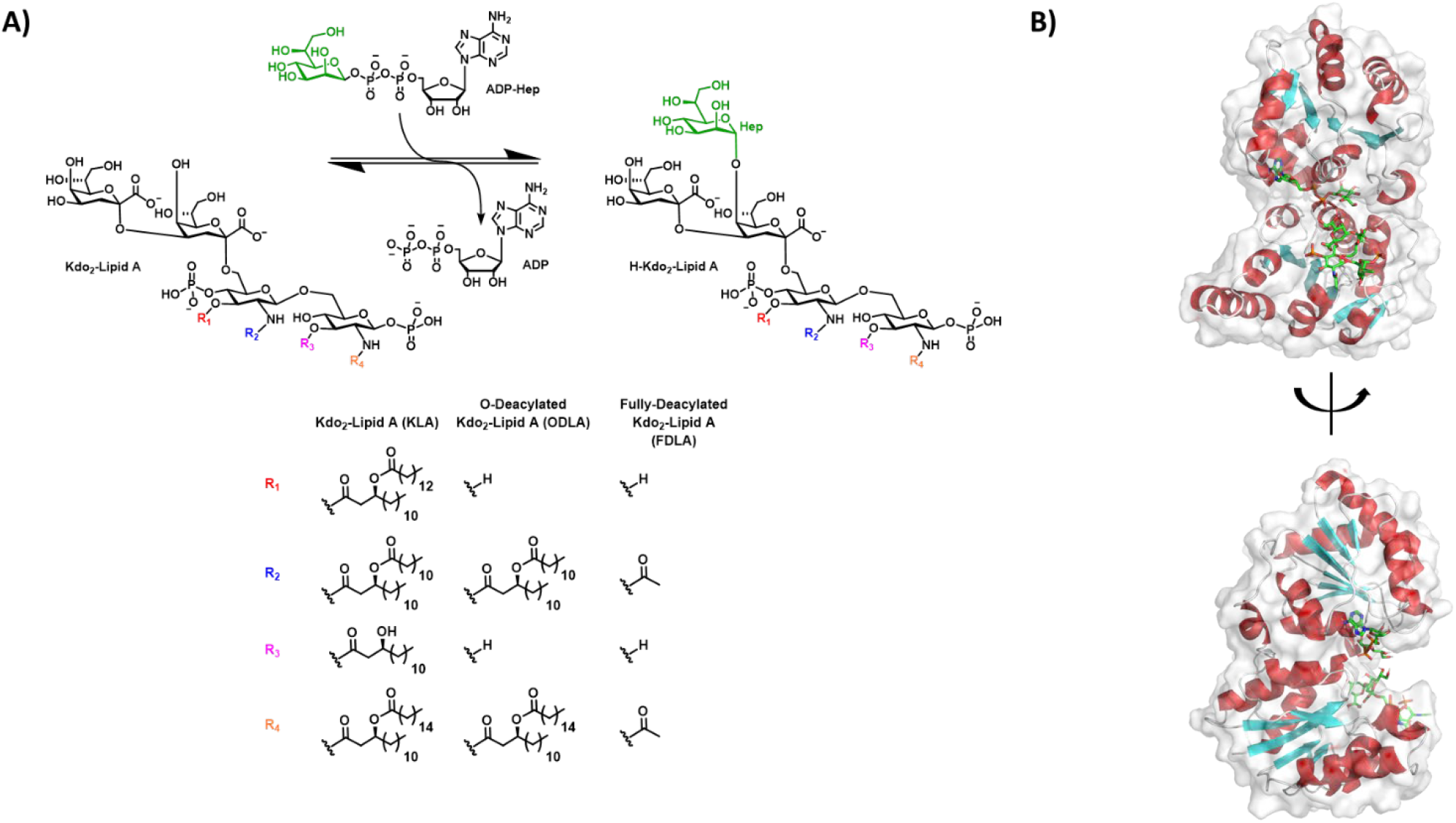
A) Reaction catalyzed by Heptosyltransferase I with deacylated derivatives used in experiments (ODLA) and molecular dynamic simulations (FDLA) highlighted based on acyl substituents. B) Fully bound substrate ternary model of Heptosyltransferase I colored by secondary structure (red: α-helix, cyan: β-strand, white: random coil).

Previous attempts at inhibiting HepI have yielded micromolar inhibitors, similar to the *K*_*M*_ values of substrate, which aren’t sufficiently potent for further development and optimization (7) (12). Grizot et. al. demonstrated a fluorine substituted ADP-Heptose analog that performed well as a competitive inhibitor of the sugar donor binding site with a mid-micromolar affinity; this compound was later used in a liganded crystal structure of the protein (PDB: 2H1H) (11). Moreau et. al. in 2008 ran the first virtual high-throughput screening with 5 million molecules to generate a small library of compounds, which were further evaluated experimentally to determine micromolar affinity for the best ligands (7). Tikad and coworkers in 2016 took a novel approach by generating a series of sugar substituted Kdo glyco-clusters and managed to achieve low micromolar inhibition with HepI; however, there seemed to be a weak bioavailability of these compounds due to their large size which violated Lipinski’s rules (13). The authors also observed aggregation of HepI with their tightest binding inhibitor *in vitro* making the overall success of their experimental findings unclear. In 2018, Nkosana et. al. synthesized monosaccharide lipid A analogs to probe the sugar acceptor binding site yielding high micromolar inhibitors of HepI. While these inhibitors had poor potency, they were the first to demonstrate that HepI could be inhibited by compounds with non-competitive inhibition mechanisms (14).

The inability to potently inhibit HepI in these endeavors suggests that alternative strategies, including seeking non- and un-competitive inhibitors that impact the protein conformational changes may be necessary to effectively inhibit it. HepI, like other GT-B enzymes of its family, has been predicted to undergo an open-to-closed transition about a hinge region, with various studies from our lab (both experimental and computational) showing that the interconversion is driven by ligand binding. For example, studies by Czyzyk et. al. which included kinetic viscosity studies as well as fluorescence stop-flow pre-steady state kinetics, showed that enzyme dynamics are partially rate limiting and play an integral role in HepI substrate binding and catalysis (15). These dynamical motions were further examined by Cote et. al. where a variety of mutations were made throughout the protein which were then examined using kinetic assays, circular dichroism spectroscopy and intrinsic tryptophan fluorescence studies. These experiments revealed amino acids responsible for ligand binding interactions necessary for initiating the open-to-closed transition (16) (17). Collectively, the data described above exposed essential information about the conformational transitions of HepI that occur in the presence of an analog of its native substrate, O-deacylated Kdo_2_-Lipid A (ODLA). Computational studies of HepI, in the presence and absence of its native substrates, helped to reveal an overall structural model for the conformational changes occurring during catalysis which can provide powerful insights for the design of HepI inhibitors (18).

Structural similarity inspection combined with experimental validation, enabled the discovery of five commercially available aminoglycosides that inhibit HepI, two of which exhibit nanomolar affinity (Figure 2, Table 1). With aminoglycoside antimicrobials previously shown to cause changes to cell membrane permeability, with a variety of possible mechanisms proposed for that effect (19), (20,21), we herein demonstrate that these compounds potently inhibit HepI. Their novel mechanism of action has been demonstrated through kinetic analyses, circular dichroism spectroscopy, intrinsic tryptophan fluorescence and molecular dynamic simulations. While these aminoglycosides resemble the tetrasaccharide core of Kdo_2_-Lipid A, we have demonstrated that while some bind to the N-terminal domain, others bind a pocket near the HepI hinge region. This binding pocket was previously identified as a potential binding pocket for the heptose portion of an ADPH analog (22) and we hypothesize that ligand binding in this pocket disrupts protein dynamics essential for catalysis.

**Figure 2:**
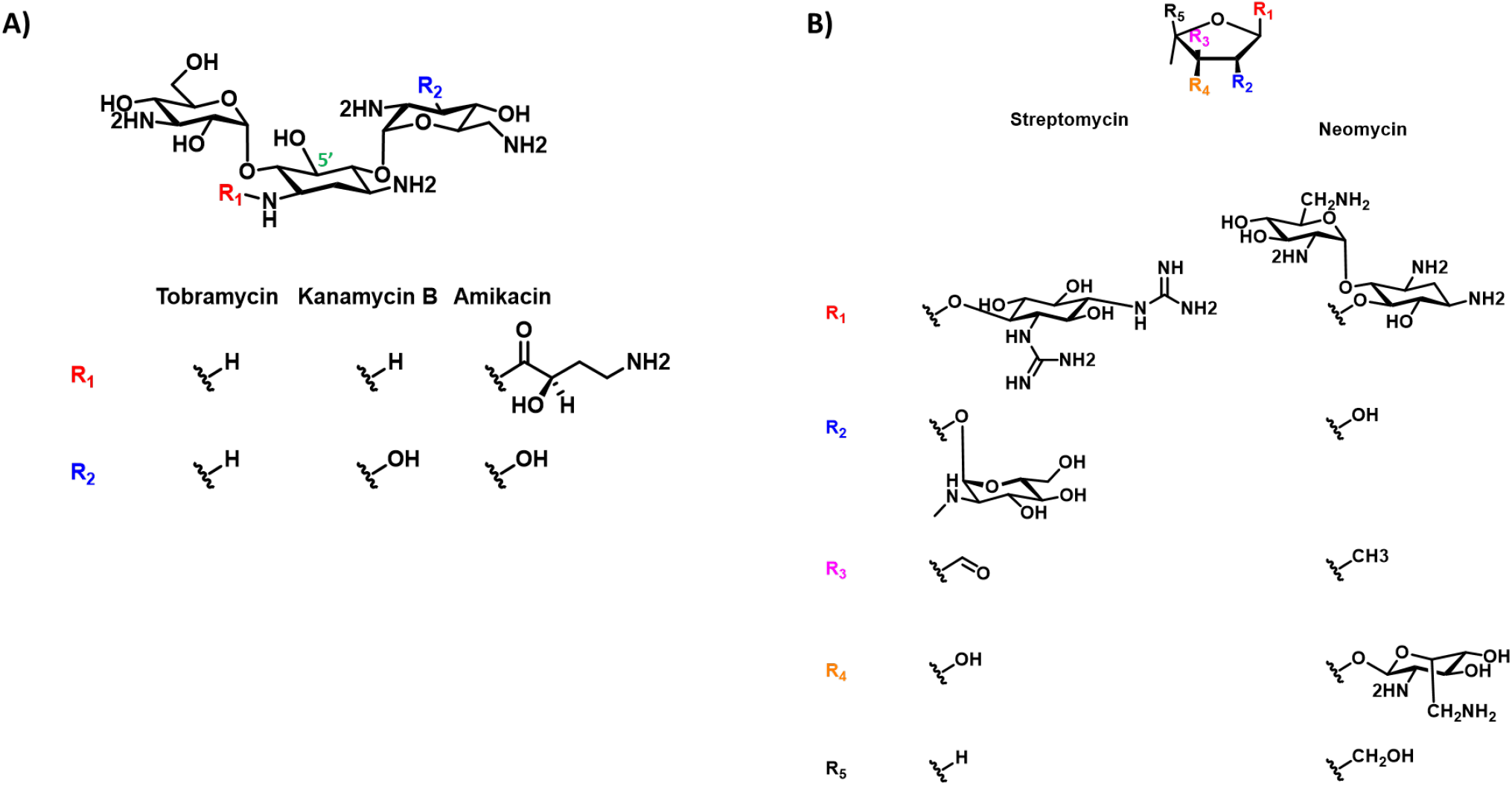
A) Trisaccharide and B) pentose scaffold of aminoglycoside inhibitors explored in this work with carbons labeled in green.

**Table 1:** A) Kinetic parameters of HepI in the presence/absence of aminoglycoside inhibitors and mutants of binding residues.

| HepI |  | Reaction Kinetics |  |  |  |  |
| --- | --- | --- | --- | --- | --- | --- |
| Inhibitor | Mutation | $k_{cat}(s^{-1})$ No inhibitor | $K_M(\mu M)$ No Inhibitor | $k_{cat}(s^{-1})$ | $K_i(\mu M)$ | Type of inhibition with ODLA |
| Amikacin | WT | $0.36 \pm 0.02$ | $2.9 \pm 0.6$ | $0.138 \pm 0.001$ | $4.258 \pm 0.631$ | N/A |
| Neomycin | | | | $0.156 \pm 0.007$ | $1.643 \pm 0.284$ | Competitive |
| Kanamycin B | | | | $0.149 \pm 0.007$ | $0.99 \pm 0.119$ | Competitive |
| Tobramycin | | | | $0.15 \pm 0.005$ | $1.125 \pm 0.212$ | Mixed competitive |
| Streptomycin | | | | $0.162 \pm 0.008$ | $0.595 \pm 0.09$ | Non-competitive |
| Tobramycin | R60A | $0.198 \pm 0.009$ | $6 \pm 1$ | $0.111 \pm 0.013$ | $3.4 \pm 0.631$ | N/A |
| Tobramycin | R120A | $0.32 \pm 0.03$ | $15 \pm 3$ | $0.829 \pm 0.044$ | $0.422 \pm 0.186$ | N/A |
| Streptomycin | R60A | $0.198 \pm 0.009$ | $6 \pm 1$ | $0.149 \pm 0.013$ | $6.254 \pm 5.427$ | N/A |
| Streptomycin | R61A | $0.36 \pm 0.01$ | $2.5 \pm 0.4$ | $0.142 \pm 0.031$ | $7.89 \pm 3.592$ | N/A |
| Streptomycin | R63A | $0.41 \pm 0.03$ | $10 \pm 2$ | $0.198 \pm 0.024$ | $6.663 \pm 0.699$ | N/A |
| Streptomycin | K64A | $0.33 \pm 0.02$ | $8 \pm 2$ | $0.117 \pm 0.009$ | $5.756 \pm 0.485$ | N/A |

## Results

### Kinetic Analysis of putative inhibitors

Inhibition constant (*K*_*i*_) values were determined for each aminoglycoside by varying inhibitor concentration at constant substrate concentration for all five putative inhibitors (Figure 3). All compounds inhibited HepI to varying degrees as shown in Table 1, exhibiting low micromolar to high nanomolar affinity which is noteworthy for a series of first generation inhibitors. Amikacin was the most ineffective inhibitor, with a *K*_*i*_ of 4.3 μM while neomycin and tobramycin exhibited slightly better *K*_*i*_ values of 1.6 and 1.1 μM, respectively. The best two inhibitors were kanamycin and streptomycin both of which displayed high nanomolar inhibition constants at 990 nM and 595 nM, respectively. Tobramycin and streptomycin were two potent inhibitors with different structural scaffolds, tobramycin having a trisaccharide structure made up of exclusively 6-membered rings, while streptomycin is a trisaccharide with a ribose ring among the 6-membered rings (Figure 2).

**Figure 3:**
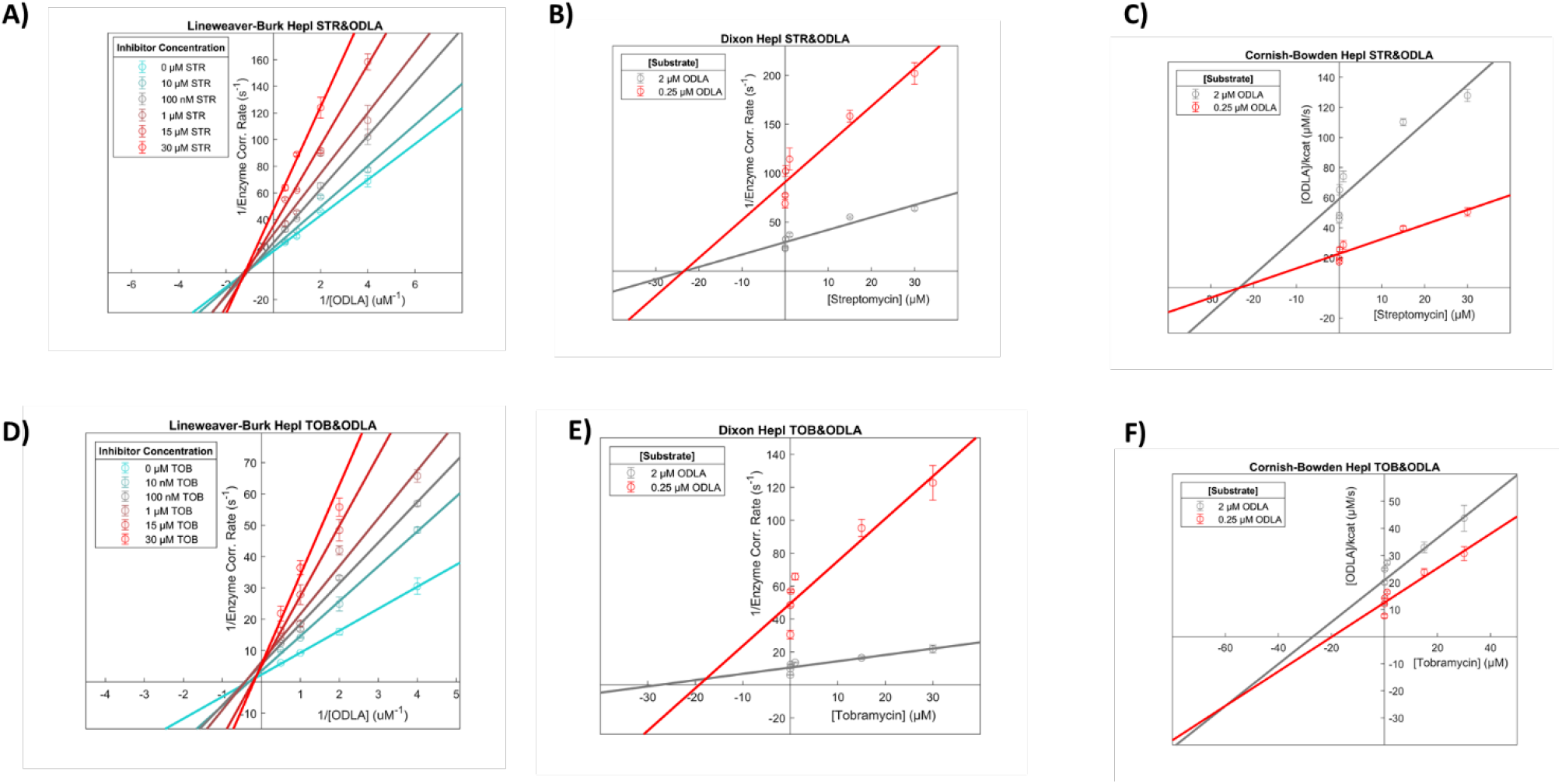
Double reciprocal plots of A-C) Streptomycin and D-F) Tobramycin in the presence of ODLA.

Further inhibition experiments were performed with the four best inhibitors, two of each structural category: neomycin, tobramycin, kanamycin, and streptomycin. These were performed at multiple substrate concentrations with varying inhibitor concentrations to determine the type of inhibition using Lineweaver-Burk, Dixon and Cornish-Bowden analyses (Figure 3A–F). By Lineweaver-Burk analysis tobramycin appears to be competitive with ODLA; however, upon careful inspection of the data, the curves do not intersect exactly at the y-axis which led to the need for additional analysis methods (Figure 3D). Reanalysis of the data with the Dixon equation yielded a plot that was again ambiguous, where competitive and non-competitive mechanisms couldn’t be distinguished because the lines intersect very close to the x-axis (Figure 3E). The Cornish-Bowden method provided the necessary clarification of the inhibition type by yielding a plot that was characteristically mixed-competitive; if the inhibitor had been purely competitive in nature, the curves would have remained parallel with the Cornish-Bowden analysis (Figure 3F). Using the mixed-model of inhibition to parse out the types of inhibition that contribute to the mixed profile of Tobramycin with ODLA, we calculated an α value greater than one which suggests a competitive/non-competitive combination. Both kanamycin and neomycin also demonstrate competitive inhibition against ODLA using the same three analysis methods (Supplemental Figure 4).

Tobramycin displays uncompetitive inhibition against ADPH, corroborated by all three analysis methods (Supplemental Figure 2D–F). This is unsurprising because ODLA and ADPH have two distinct binding sites and tobramycin is unlikely to compete with both simultaneously due to its size. Thus, it is possible that tobramycin may bind to a region of the protein that is only accessible upon structural rearrangements after ADPH binding. The competition of tobramycin with ODLA was further investigated by examining changes in *K*_*i*_ values of tobramycin using mutant HepI constructs previously identified to alter the binding of ODLA (17) (Supplemental Figure 5E–F, Table 1). Specifically, mutation of the positively charged residues Arginine 60 and 120 to Alanine resulted in alterations of tobramycin inhibition of HepI (Table 1). The HepI R60A mutant exhibited a *K*_*i*_ approximately 10-fold weaker relative to the WT, while the R120A demonstrated a *K*_*i*_ ~2-fold tighter than the WT.

Streptomycin exhibited non-competitive inhibition against both ODLA and ADPH, which was unambiguously evident with all three analysis methods (Figure 3A–C, Supplemental Figure 3D–F). Streptomycin binding to the enzyme was also investigated by way of *K*_*i*_ analysis using mutant forms of HepI including: R60A, R61A, R63A, and K64A (Table 1, Supplemental Figure 5A-D). Varying concentrations of Streptomycin with all four mutant forms increased the *K*_*i*_ values an order of magnitude suggesting that these residues were not just proximal to the Streptomycin binding site but are also mediating the binding of the compound.

### Circular Dichroism Thermal Analysis

Since kinetic analyses indicate that tobramycin is partially competitive with ODLA for binding, circular dichroism studies were performed to assess if tobramycin disrupts the previously observed ODLA-induced thermal stabilization of HepI (17). Previous studies of HepI WT revealed that the apo protein melts at around 37°C and unfolds ~80% from the fully folded, whereas in the presence of ODLA, HepI shows an increase in thermal stability and melts above 95°C (Figure 4, Supplemental Figure 6A-B). The addition of ADPH to WT HepI does not induce any changes in the melting temperature but does destabilize the protein to allow it to unfold fully (~100%). Unsurprisingly, HepI in the presence of the heptosylated ODLA (ODHLA) product behaves like HepI and ODLA with no observable melting event below 95°C and HepI with ADP melts similarly to HepI with ADPH (Supplemental Figure 8).

**Figure 4:**
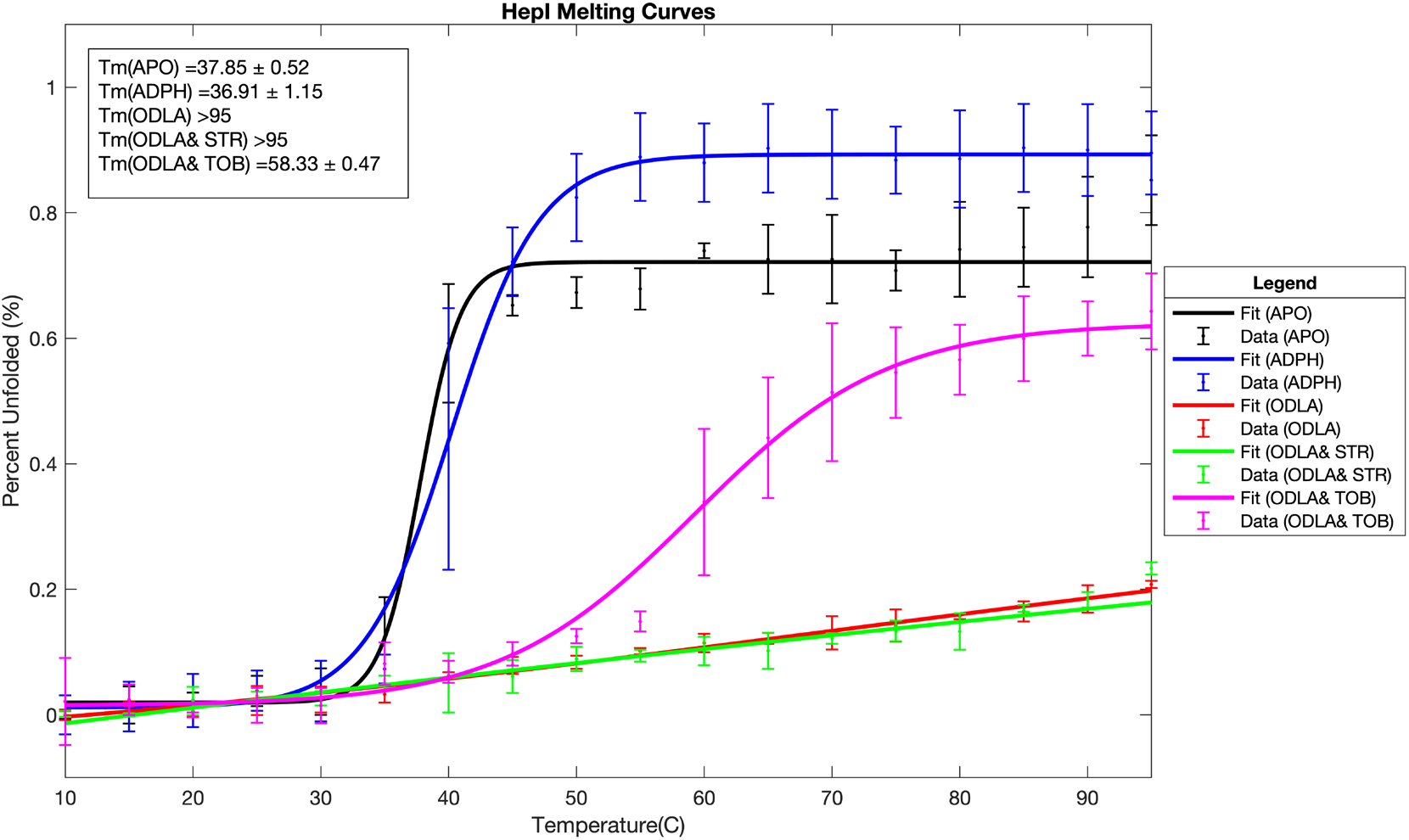
HepI melt curves in the presence of substates and aminoglycoside inhibitors.

The HepI•tobramycin complex has a Tm of ~37°C with a 20% increase in unfolding as compared to apo (Figure 3, Supplemental Figure 7D), suggesting an induced destabilization of the enzyme. The addition of tobramycin to the HepI•ODLA complex disrupts the ODLA-induced thermal stabilization resulting in a protein complex that melts at around ~58°C and unfolds to about 70% (Figure 4, Supplemental Figure 6C,7-8). HepI with ADPH and tobramycin showed no changes from HepI with ADPH alone. These findings are corroborated by the kinetic analysis of tobramycin being a mixed-competitive inhibitor against ODLA. A CD melt analysis of the HepI•tobramycin complex with each of the products behaves like their substrate counterparts.

HepI with streptomycin melts at 37°C with a ~20% increase in unfolded protein in comparison to apo and no discernable difference in secondary structure (Supplemental Figure 6C, Supplemental Figure 7A). Streptomycin in the presence of either substrate or product appears no different from substrate or product alone (Supplemental Figure 7C-D, Supplemental Figure 8B). This aligns well with the kinetics data that suggests that streptomycin is non-competitive with both substrates. The inhibitor is free to bind to any form of HepI (E, E•S, E•S_2_, E•P_2_, E•P) and interrupt chemistry.

### Intrinsic Tryptophan Fluorescence

Characteristic behavior of HepI in fluorescence studies have shown that formation of the HepI•ODLA complex results in a 6 nm blue shift in the HepI Tryptophan (Trp) emission spectra (Figure 5A). This method was a good reporter of global protein rearrangements that caused Trps to become less solvent exposed due to ODLA binding. If tobramycin disrupted the thermal stability of HepI with ODLA it was then hypothesized that it would truncate the blue shift however, it did not (Figure 5B, Supplemental Figure 10B). HepI with tobramycin did not alter the λ_max_ and when adding tobramycin to the HepI•ODLA complex, a 6 nm blue shift was still observed. Furthermore, the same phenomenon was observed with streptomycin where the substrate induced blueshift was unperturbed. HepI•tobramycin and HepI•streptomycin complexes had no changes to the emission spectra with respect to HepI apo (Figure 5, Supplemental Figure 10).

**Figure 5:**
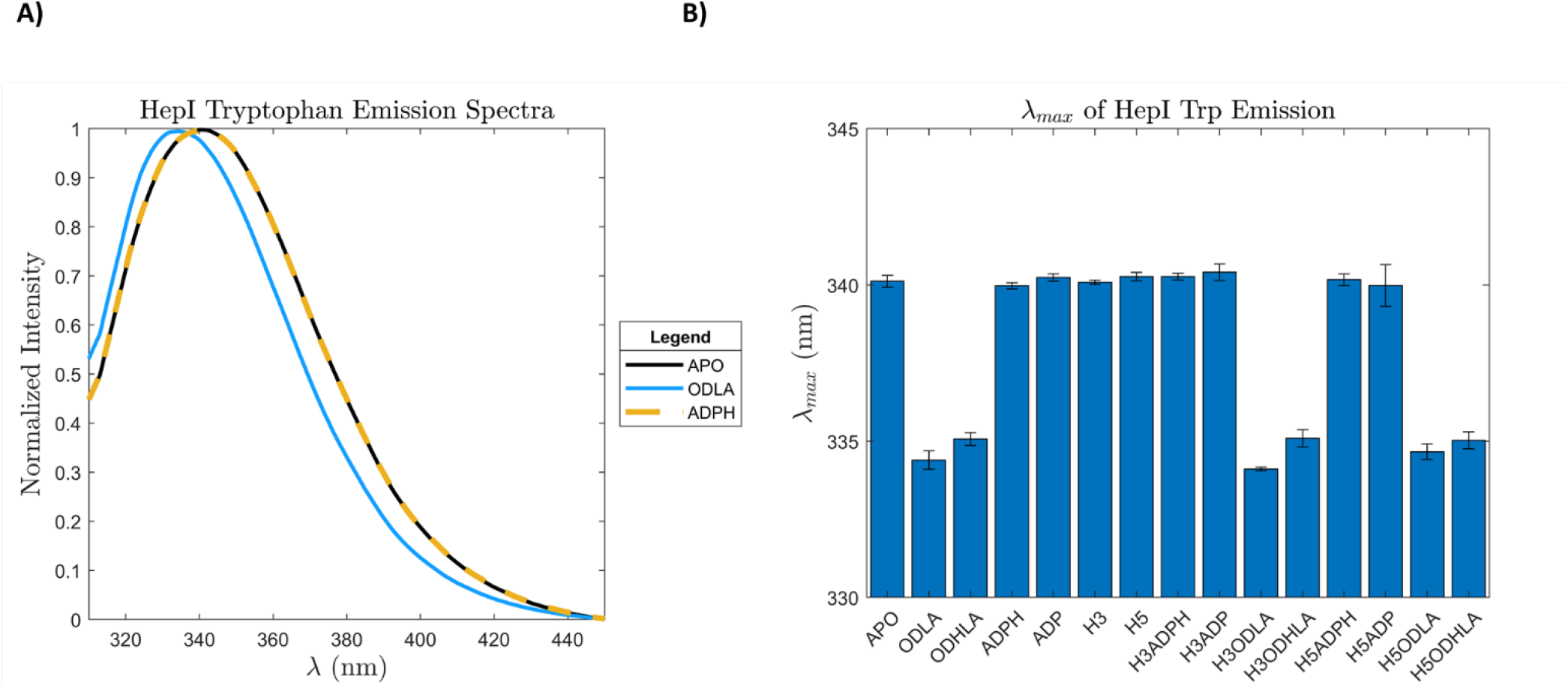
HepI tryptophan fluorescence emission A) spectra and B) λ_max_ in the presence/absence of substrates, products and inhibitors.

### Inhibitor Docking and Protein-Inhibitor Interactions Dynamics

To gain a better understanding of the possible interactions between HepI and these aminoglycoside inhibitors, we turned to docking and molecular dynamics simulations guided by our experimental evidence. While experiments were performed with ODLA, a fully deacylated analog of Kdo_2_-Lipid A (FDLA) was used for all computational studies (Figure 1A). Docking experiments showed binding poses of tobramycin to the HepI•ADPH•FDLA and HepI•ADPH complex which revealed a pocket that is between the FDLA and the heptose portion of the ADPH (Figure 6C). Furthermore, streptomycin docked to a pocket that is below the adenine ring of ADPH in the HepI•ADPH•FDLA complex (Figure 6B). Binding free energies for tobramycin and streptomycin are outlined in supplemental table 3.

**Figure 6:**
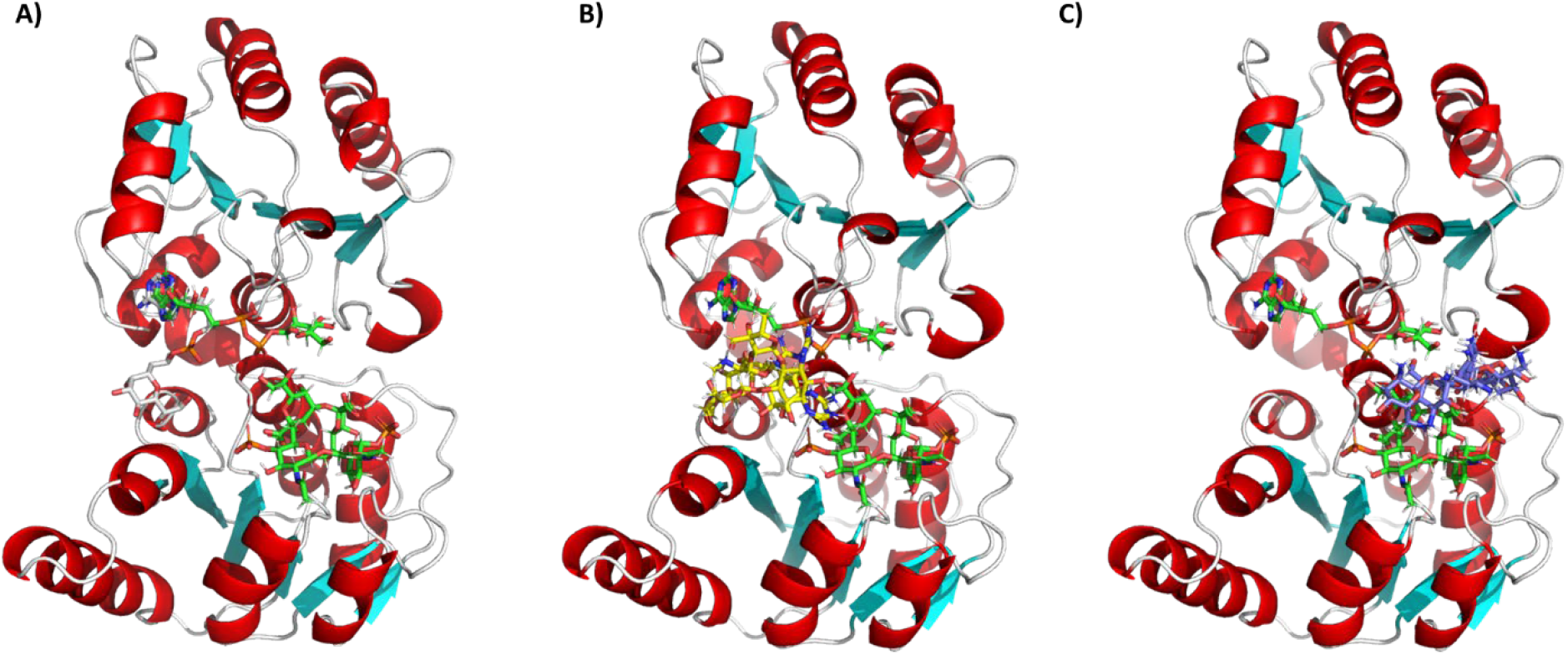
(A) Model of HepI•ADPHep •FDLA complex with the native substrates (green) and the original pseudo phosphonate derivative (white) and (B) the top two binding poses for Streptomycin (yellow) and (C) Tobramycin (blue).

Simulations of HepI, HepI•ADPH•FDLA, HepI•ADP•FDHLA, HepI•ADPH•FDLA•Tobramycin, HepI•ADPH•Tobramycin, HepI•ADPH•FDLA•Streptomycin, HepI•ADP•FDHLA•Tobramycin, HepI•ADP•FDHLA•Streptomycin were performed for 100 nanoseconds, each (Supplemental Table 1). Stability of each system was determined by the average backbone root mean square deviation (RMSD) and the C-alpha radius of gyration (CαR_gyr_). The average RMSD of all the systems coalesce to a value under 2 Å (Supplemental Figures 11A-14A, Supplemental Table 2). As demonstrated in supplemental table 2, the average RMSD for HepI substrate ternary complex, in the presence of tobramycin, or streptomycin are 1.84 ± 0.23 Å, 1.88 ± 0.30 Å, 1.84 ± 0.25 Å, respectively. Meanwhile, the average RMSD for HepI product complex, in the presence of tobramycin and streptomycin are 1.62 ± 0.26 Å, 1.59 ± 0.33 Å, and 1.62 ± 0.26 Å, respectively. The R_gyr_ for the HepI ternary complex, in the presence of tobramycin and streptomycin are 21.13 ± 0.16 Å, 21.23 ± 0.12 Å, and 21.28 ± 0.16 Å, respectively. In the HepI product ternary complex and in the presence of tobramycin and streptomycin, the average R_gyr_ is 21.09 ± 0.16 Å, 21.32 ± 0.90 Å, and 21.16 ± 0.14 Å, respectively.

Neither metrics display large structural deviation over time, indicative of a tightly bound stabilized closed complex. Most residues in the protein display very little fluctuations and only a handful of residues exhibit fluctuations greater than 1.5 Å as determined by the C-alpha root mean square fluctuation (C_α_RMSF) (supplemental table 2, supplemental Figure 11A/B-14A/B), as is consistent with prior studies of the HepI apo and substrate complexes (22). The HepI•ADPH•FDLA complex has residues with greater than 1.5 Å fluctuations in the N_3_ loop (63–69), C_1_ loop (207,209), and C_α6_ (300,301,317-324). Similarly, the HepI•ADPH•FDLA•tobramycin displays nearly identical regions of greater than 1.5 Å fluctuations including residues in N_α3_ (63-69,71), C_1_ loop (187), C_α1_ (207,209), C_α2_ (231), C_5_ loop (280), C_α5_ (284–286), and C_α6_ (300,301,317-324). Finally, the HepI•ADPH•FDLA•streptomycin complex, residues with greater than 1.5 Å fluctuations include N_α3_ (62-69,72), N_6_ loop (135-136), C_α1_ (207), C_5_ loop (280), C_α5_ (285-288), and C_α6_ (300,317-324). However, in the presence of the products, dynamic residues include N_α3_ (60-61,63-73,84), N_6_ loop (137), C_5/6_ loop(289), and C_α6_ (317-320). Furthermore, tobramycin in the presence of products, HepI•ADP•FDHLA•tobramycin, has nearly identical dynamic residues which include N_α3_ (61-77,84), N_α6_ (136-137,156-158), C_α5_ (286), C_5/6_ loop (291-292), and C_α6_ (318-320). On the other hand, streptomycin in the presence of the products, HepI•ADP•FDHLA•streptomycin, has fewer dynamic residues that include N_α3_ (63-73) and C_α6_ (318-320). The RMSF is useful for highlighting residues with large fluctuations but can be difficult for discerning small differences in fluctuations however, taking the difference of RMSFs provides relative fluctuations that may have otherwise gone unnoticed. The ΔRMSF for the HepI ternary complex in the presence of tobramycin or streptomycin relative to the HepI ternary complex in the absence of inhibitors displays changes in Nα4, N_α6_, and C_α5_ (supplemental Figure 11C-14C, Supplemental Table 2).

As stated previously, streptomycin occupies a pocket below the adenine ring of the ADPH and adjacent to the FDLA (Figure 6B), somewhat near the linker that connects the N− and C-terminal Rossmann-like domains. Streptomycin maintains a relatively stable position throughout the 100 ns trajectory. A ligand interaction diagram for the last frame of a representative simulation reveals only one potential hydrogen bonding partner, S10 (Figure 7A). Upon further analysis, residues with an average minimum distance of less than 3.5 Å to streptomycin include ones in N_α1_(9–11), N_2_ loop (38,40,41), N_α3_(60,63,64), C_α1_(188-190), and C_α2_(218-219,221-222) (Figure 7C). The average minimum distance of streptomycin to ADPH or FDLA is less than 2 Å. The streptomycin C1’-6’ carbons are closest to the second KDO sugar on the C4 phosphate side. In contrast, tobramycin binds to a pocket sandwiched between the heptose moiety of ADPH and the FDLA (Figure 6C). A ligand interaction diagram for tobramycin reveals an electrostatic attraction between a charged primary amine on tobramycin and the putative catalytic residue D13 (Figure 7B). Residues that have an average minimum distance of less than 3.5 Å to tobramycin include N_α1_(13), Nα5(120), N_α6_(136-137,140,143), C_α1_(188-189,191-192), and C_α5_(281-283,286-287) (Figure 7D). In this case, tobramycin is stacked directly above FDLA and has average minimum distance less than 2 Å, whereas the ADPH is greater than 2 Å (Figure 7D). Tobramycin C1”-6” sugar stacks on top of the terminal KDO sugar, whereas the C1-6 and C1’-6’ sugar/psuedosugar stack with the heptose portion of ADPH.

**Figure 7:**
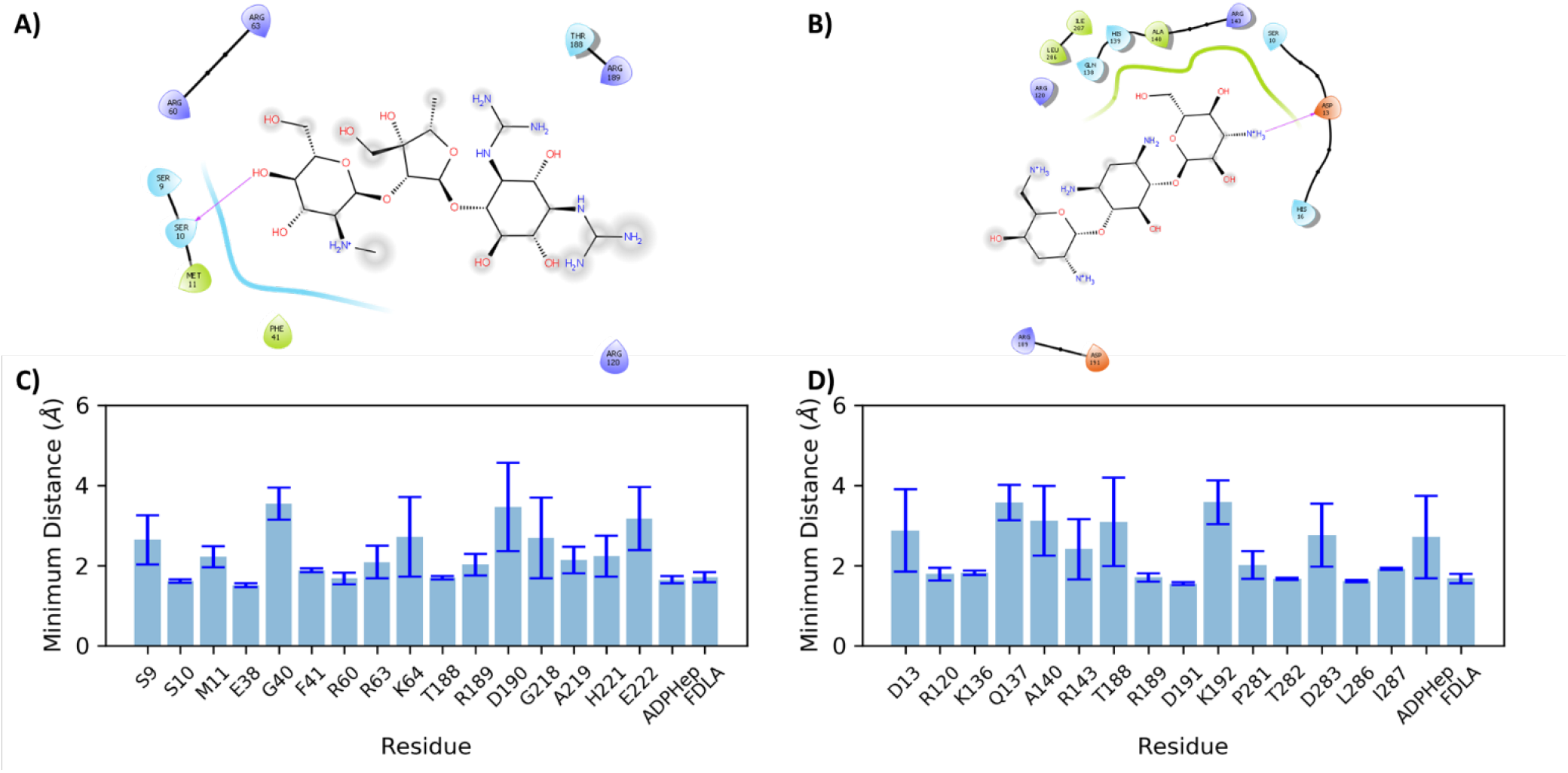
(A) Streptomycin Ligand Interaction Diagram (B) Tobramycin Ligand Interaction Diagram (C) Residues with minimum distance <3.5 Å from Streptomycin and (D) Residues with minimum distance <3.5 Å from Tobramycin

Comparing the average pairwise distance between atoms of the substrates and inhibitors reveals electrostatic interactions between the phosphates of the substrates and the amines of the inhibitors. The average distance between the charged secondary amine on the C2’’ carbon of streptomycin and the alpha phosphate oxygens of ADPH is 3.7 ± 1.0 Å (Supplemental Figure 15A,C). The primary amine of the streptomycin guanidinium group on the C8’ carbon is 7.6 ± 0.9 Å from the beta phosphate oxygens of ADPH. Relative to C4 GlcNAc phosphate oxygens of the FDLA, the streptomycin primary and secondary amines on the C7’ carbon are 6.8 ± 1.1 Å and 6.8 ± 1.5 Å, respectively. The tobramycin charged primary amine on the C6” carbon is 4.5 ± 0.9 Å from the C1 GlcNAc phosphate oxygens of FDLA. In the binary complex, the tobramycin primary amine is 3.3 ± 0.3 Å from the beta phosphate oxygens of ADPH.

## Discussion

The pursuit of a competitive inhibitor for HepI has been driven by the need to impede *in vivo* LPS biosynthesis for antimicrobial therapeutic design. Additionally, since Kdo_2_-Lipid A induces a global conformational change in HepI, inhibiting HepI could also lead to catalytically unproductive conformational changes in the enzyme, thereby reducing catalysis and subsequently the LPS coverage on the bacterial surface. The structural resemblance of the tetrasaccharide core of Kdo_2_-Lipid A to a collection of aminoglycosides has led to the discovery of potent inhibitors for HepI.

The preliminary evaluation of the aminoglycoside library revealed two first-in-class, nanomolar inhibitors of HepI, streptomycin and kanamycin b. Kinetic and biophysical characterization of these inhibitors has revealed their mechanisms of action and binding sites. These inhibitors were grouped for evaluation based on their structural similarities to one another. Streptomycin and neomycin both have ribose rings mediating glycosidic bonds to 6-membered rings. Kanamycin b and tobramycin are both trisaccharides composed entirely of 6 membered rings. Kinetic analysis of inhibition with tobramycin revealed a mixed-competitive behavior against ODLA and uncompetitive inhibition against ADPH, corroborated by all three kinetic analysis methods, described above. Kanamycin b showed competitive inhibition with ODLA; as a structural analog of tobramycin, kanamycin b that only differs by one hydroxyl group, we hypothesized that kanamycin’s inhibitory character would resemble that of tobramycin with ADPH.

Streptomycin was determined to act by a non-competitive mechanism with both native substrates, as described above. We hypothesized that the ribose ring would create a structural similarity between neomycin and streptomycin since both were nanomolar affinity inhibitors. Interestingly, neomycin exhibited a purely nanomolar competitive mechanism for inhibiting ODLA binding to HepI, as corroborated by our kinetic analyses. Neomycin being a tetrasaccharide may adopt a binding configuration that resembles the sugar acceptor substrate (Kdo_2_-Lipid A) more effectively than tobramycin and kanamycin b, which are both trisaccharides. These data suggest that a tetrasaccharide structural scaffold is more important than sugar species in inhibition competitively with ODLA (Figure 2, Table 1). When neomycin is superimposed on the structure of FDLA, the hexoses/psuedohexos of neomycin mimic a similar configuration of the two Glucosamines and terminal KDO of FDLA (Supplemental Figure 17).

Docking and simulations of HepI as a substrate or product ternary complex in the presence of tobramycin or streptomycin demonstrated interesting variations in binding pockets and their effects on the conformational change required prior to catalysis which allowed the generation of models for the inhibition by these two classes of inhibitors. Streptomycin occupies a binding site that is near the hinge region between the two Rossman-like domains (Figure 6B). This site was previously identified through as the binding site of the heptose moiety of a phosphonate derivative of ADPH in a previously solved HepI pseudo-ternary substrate complex (Figure 6A) (23). Additionally, in our previous simulations of HepI with ADPH (data not shown), we observed the heptose moiety flip into this pocket over the course of a 50ns simulation. This pocket seems to accommodate sugar binding and is therefore plausible non-competitive, allosteric inhibition pocket.

Previous studies revealed that the HepI N_3_ loop, adjacent to this pocket, is highly dynamic and contains multiple residues that are involved in binding of FDLA through electrostatic interactions with redundant positively charged residues. The presence of either inhibitor does not perturb the observed molecular dynamics of this region. Streptomycin instead interacts with the phosphates on the ADP-facing side of the FDLA, but rather than outcompete the positively charged arginine/lysines, the multivalent negative charges of the phosphate can accommodate interacting with the enzyme, while also interacting with streptomycin. Furthermore, this distance between the phosphate of FDLA and the positively charged amines on streptomycin are so far apart that streptomycin will have a weak influence on the FDLA from that phosphate at best (Supplemental Figure 15A,C). The guanidinium group on streptomycin interacts with the alpha phosphates of ADPH to further provide another anchor in that site, but streptomycin is within hydrogen bonding distance to several negatively charged residues (E38, D190, E222) that could replace this interaction (Figure 7C). Therefore, streptomycin could bind to the enzyme in the absence of these two ligands by utilizing these other negatively charged residues while staying far away enough from the crucial FDLA binding residues to constitute its place as a noncompetitive inhibitor for both substrates.

Alternatively, tobramycin binds to a cleft between the heptose and the FDLA (Figure 6C). Experimental evidence suggests tobramycin acts as mixed (competitive/noncompetitive) inhibitor with FDLA, and an uncompetitive inhibitor of ADPH. This binding site model provides a convenient location for tobramycin to interact with ADPH in an un-competitive manner while simultaneously positioning it to replace FDLA in its absence in a competitive fashion. Additionally, binding in this pocket would allow tobramycin to act as a dynamics disruptor of the open-to-closed conformational transition which would be non-competitive with ODLA or FDLA binding. Tobramycin has a positively charged amine near the FDLA phosphate with two other positively charged residues that are within 3.5 Å (Arg120, Lys136) (Figure 7D, Supplemental Figure 15B-C). This will lead to an unfavorable electrostatic repulsion between these positively charged residues and tobramycin. Alanine mutation of Arg120 leads to the lowering of the inhibition constant of tobramycin, which supports this hypothesis (Table 1). Alanine mutation of K136 may be able to achieve this affect without perturbing the binding of FDLA. Alternatively, the reduction in charge of tobramycin through derivatization may lead to tighter binding inhibitor with similar properties.

In our previous molecular dynamics simulations, we observed a conformational rearrangement of HepI in the presence of its products on the nanosecond timescale reminiscent of that which is expected from a GT-B prior to catalysis. Since this rearrangement occurs prior to catalysis, we used this as a proxy to computationally explore whether these inhibitors affect the ability of the enzyme to undergo the open-to-closed transition. Specifically, we calculated the distribution of the center of mass for the enzyme in the presence of each inhibitor and compared to the uninhibited enzyme. (22) Based on the computationally determined center of mass distributions, streptomycin has no effect on this rearrangement, whereas tobramycin hinders the closure that can be observed over these short time periods (Figure 8).

**Figure 8:**
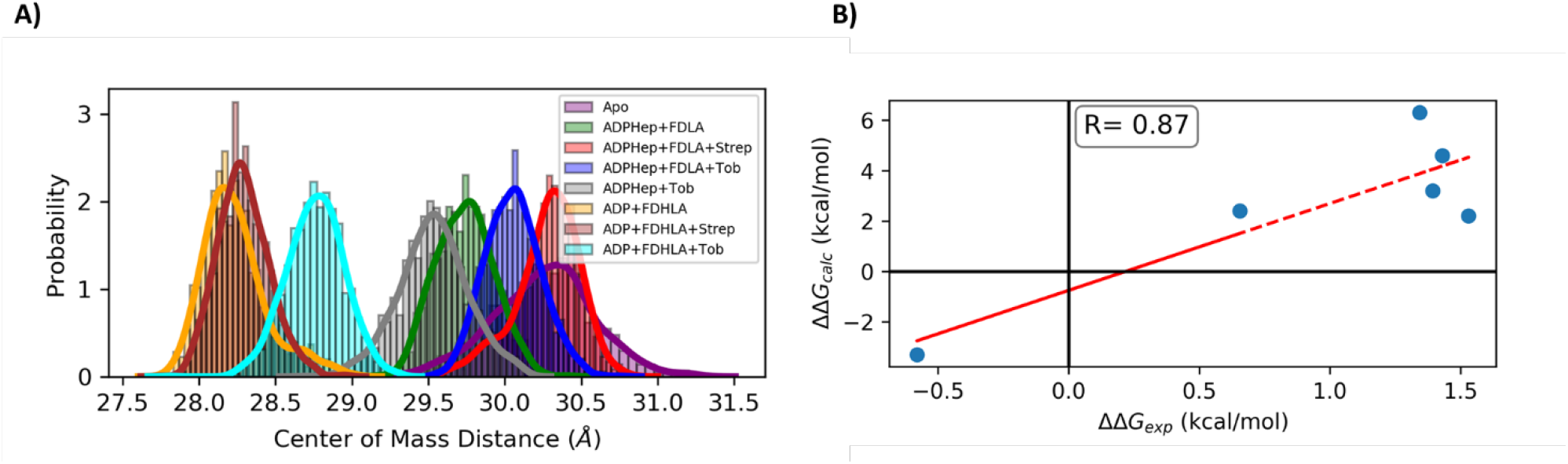
(A) Probability distribution of center of mass distance between N and C termini of HepI in the apo (purple), substrate (green), substrates with Streptomycin (red), substrates with Tobramycin (blue), HepI binary with Tobramycin (grey), products (orange), products with Streptomycin (brown), and products with Tobramycin (cyan). (B) Binding free energy differences of Tobramycin/Streptomycin to HepI WT relative to ARG/LYS mutants.

The biophysical characterization of HepI inhibition corroborates mechanistic behavior revealed in these molecular dynamics simulations. Previous studies that investigated the residues in HepI integral to the binding of ODLA showed a series of positively charged residues found on loops flanking either side of the ODLA binding site creating vital electrostatic interactions to the substrate. These loops rearrange to accommodate the substrate by becoming more alpha helical. Proximal to these positively charged residues are W62 &116, previously shown to make the largest contribution to the blue shift seen with ODLA binding. Uninhibited HepI exhibits a 6 nm blue shift of the tryptophan λ_max_ upon binding of ODLA, which can serve as an indirect reporter for binding and protein rearrangement caused by ODLA (Figure 5A) (17). This spectral shift is attributed in part to changes in the hydrophobicity of the residues Trp62 and Trp116, which are on loops involved in the binding of ODLA. Neither of these inhibitors affect the steady-state tryptophan fluorescence blue shift, which is consistent with the simulations above (Figure 5B). Since there are no changes to the blue shift due to the addition of inhibitor, and we hypothesize that the loops enriched with positive charges are still able to rearrange to accommodate ODLA.

Over the course of the simulation, streptomycin does not affect the secondary structure of the protein, whereas tobramycin influences this secondary structure and the ability of HepI to close (reduced diameter of the enzyme). Therefore, these inhibitors mostly leave the HepI•ODLA complex formation unaffected, yet they have different effects on the global conformational change induced by ODLA binding. Examination of the impact of these inhibitors on the circular dichroism studies and thermal stability of HepI•ODLA complex similarly recapitulate these differences in inhibition mechanism. While the overall CD spectra of the enzyme-substrates complexes at 10°C in the presence and absence of inhibitors were nearly identical, notable differences in the overall thermal stability between these inhibitors were observed. Where a competitive inhibitor would completely disrupt the binding of ODLA to its binding site, tobramycin partially competes for binding to the ODLA site only causing partial destabilization of the HepI•ODLA complex, a >30 °C difference in melting temperature. This affect is not observed for streptomycin, consistent with its non-competitive mechanism (Figure 4).

Additionally, thermal melt experiments in the absence of ODLA showed that the HepI•streptomycin complex destabilized the enzyme evident by an increase of about 15-20% of the unfolding in comparison to the apo with no changes in Tm. This protein destablization was also observed in HepI•ADPH, HepI•ADP, HepI•tobramycin, HepI•tobramycin•ADP/H, HepI•streptomycin•ADP/H complexes (Supplemental Figures 7A,C). Based on our experimental and computational findings, thus far, we hypothesize that this phenomenon appears due to a disruption in intrinsic electrostatic interactions within the protein that prevent it from fully unfolding in the apo form which is facilitated by the excess charges introduced by these respective ligands. In the case of a conformational change like that of HepI, the burial of charges has a high desolvation penalty and proteins overcome this barrier by neutralizing through charge pairing or protonation/deprotonation events.(24) In terms of stability, paired charges increase the stability of a protein due to the strength of electrostatic interactions. We therefore hypothesize that the paired positive charges introduced by streptomycin near the N_3_ loop have no net effect on the stabilization that results from the HepI and ODLA complex or the desolvation barrier for closing. However, the excess positive charge introduced by tobramycin into the N_5_ loop region effects the HepI•ODLA complex by increasing the desolvation barrier and preventing the full closure while also disrupting the stabilization effect of the neutralized charges through electrostatic repulsion.

## Conclusions

This small library of compounds are all aminoglycosides; a potent class of antimicrobial drugs used alone or as a combinatorial therapy against aerobic Gram-negative and a select few Gram-positive bacterial infections. Previous studies that have investigated the aminoglycoside antimicrobial activity conclude that the primary mode of action is binding to the 30s ribosome and the mistranslation of integral membrane proteins. However, it is also shown in previous studies that there are non-native “pores” that rapidly develop on the bacterial membrane which is attributed to alterations in protein translation and insertion into the membrane. We note that the rate of translation and installation of membrane proteins is far slower than the rate of LPS biosynthesis, and while the disruption of LPS biosynthesis was not taken into consideration when these findings were published, due to lack of discovery, we have an alternative hypothesis as to the mechanism of membrane destablization that occurs when treating bacteria with aminoglycosides. As has been observed in other studies, the disruption of LPS biosynthesis resulting from HepI inhibition causes a reduction in the extracellular polymeric substances embedded on the outer leaflet of the bacteria which could cause the inevitable “pores” observed on the bacterial surface (often described as a deep rough phenotype). With data given above that demonstrates the *in vitro* inhibition of HepI by aminoglycosides, we hypothesize that at least part of the mechanism of action of these antimicrobials is interruption of LPS biosynthesis.

## Materials and Methods

### Materials and equipment

*E. coli* strains DH5α and BL-21-AI were obtained from Invitrogen (Carlsbad, CA). Isopropyl β-d-1-thiogalactopyranoside (IPTG) was obtained from Gold Bio Technologies (St. Louis, MO). Amicon Ultra-15 (30 kDa, 10 kDa and 3 kDa Molecular Weight Cut-Off (MWCO)) centrifugal concentrators, Sartorius Vivaspin 20 (10 kDa MWCO) centrifugal concentrators, tryptone and yeast extract were obtained from ThermoFisher Scientific (Pittsburg, PA). Sodium chloride, sodium hydroxide, ampicillin (Amp), tetracycline (Tet), tobramycin (Tob), streptomycin (Str), amikacin, neomycin, kanamycin b, HEPES, imidazole, ethylenediaminetetraacetic acid (EDTA), cobalt sulfate, and L-arabinose were acquired from Sigma (St. Louis, MO). Bio-Scale Mini Bio-Gel P-6 desalting cartridge was obtained from Bio-Rad (Hercules, CA). All UV-Vis measurements were taken using Cary 100 Bio UV-Vis from Agilent (Santa Clara, CA). Circular dichroism spectra were measured using a Jasco J-810 spectropolarimeter (Easton, MD). Fluorescence spectra were measured using Fluoromax-2 from Horiba Scientific (Edison, NJ). All cells were lysed using a EmulsiFlex-C5 homogenizer manufactured by Avestin Inc. (Ottawa, ON). Toyopearl AF-Chelate-640 resin was obtained from Tosoh Biosciences (Grove City, OH). All ESI-MS spectra were collected using a Thermo Scientific (Waltham, MA) ESI spectrometer.

### Substrate isolation and deacylation

ADPH and Kdo_2_-Lipid A isolations have been previously reported. ADPH and Kdo_2_-Lipid A were extracted from *E. coli* WBB06 cells (HepI and HepII knockout *E. coli* strain). Overnight cultures were started by inoculating 10 mL of LB-Tet with WBB06 cells from a glycerol stock stored at −80 °C. LB-Tet media (8 L) was inoculated (1 mL of overnight growth per 2 L of media) and allowed to grow at 37 °C to an OD_600_ of 1 (approximately 12 hrs), then centrifuged for 10 min at 5,000 rpm to pellet the cells. ADPH was extracted by adding 80 mL of 50% ethanol to the pellet and stirred on ice for 2 hrs. In 30 mL Nalgene tubes, cells were centrifuged for 20 min at 10,000 rpm and supernatant was saved and the ethanol was removed by vacuum on ice. The crude extract was ultracentrifuged for 1 hr at 40,000 rpm and filtered successively using Amicon Ultra-15 30kDa, 10kDa and 3kDa centrifugal filters. Finally, the flow through was placed over a 64 mL DEAE column using a triethylamine bicarbonate (pH=8) gradient from 1-500 mM to purify, crude ADPH and water were also brought to a pH of 8. Approximately 500 μL of each fraction were lyophilized and ESI-mass spectrometry was used to determine fractions that contained pure ADPH by observation of a peak at (m/z^−1^ =619) in a 50:50 acetonitrile:water solution. Fractions confirmed to contain ADPH were pooled and lyophilized successively to remove traces of triethylamine.

Kdo_2_-Lipid A was extracted from 8 L of frozen or fresh WBB06 cells grown as described for ADPH extraction. The cells were resuspended in 80 mL of water and the mixture was divided into 30 mL Kimble glass tubes (10 mL per tube) and centrifuged for 10 mins at 5,000 rpm. The supernatant was discarded and cells were washed with 160 mL ethanol followed by 160 mL of acetone twice and finally 160 mL of diethyl ether by cellular resuspension in each of the solvents and centrifugation to discard the supernatant before the next wash. Cells were then left to dry in the hood at room temperature 1hr-overnight in a large weigh boat. Kdo_2_-Lipid A was then extracted from the dried down cells by pulverizing the cells into a fine powder and adding 20 mL of a solution per tube of 2:5:8 phenol, chloroform, and petroleum ether to 1 gram of cells per 30 mL tube. The mixture was vortexed for 3-5 mins, left on a nutator for 10 mins, then centrifuged for 5,000 rpm for 10 min. Supernatant was gravity filtered through filter paper and the extraction was repeated. The diethyl ether and chloroform was removed in vacuo from the supernatant and a mixture of 75 mL acetone, 15 mL diethyl ether and 5 drops of water was added to the solution and left to sit for 45 mins-overnight to precipitate the Kdo_2_-Lipid A. The solution was centrifuged at 5,000 rpm for 10 min in 30 mL glass tubes and supernatant was discarded. Pellets were washed a minimum of 3 times with ~1 mL 80% phenol and diethyl ether each separately; solutions were centrifuged in between washes and supernatant was removed prior to next wash. The pellets will dramatically diminish over the course of the washes and become more colorless. The dried pellets were then dissolved in 0.5% triethylamine aqueous solution and flash frozen for lyophilization (yield ~50 mg).

Synthesis of ODLA has previously been reported (25). Briefly, o-deacylation of Kdo_2_-Lipid A was done by refluxing a mixture of 5 mL hydrazine to 50 mg of extracted Kdo_2_-Lipid A for 1 hour at 37 °C in a round-bottom flask with stirring (1 mL of hydrazine for every 10 mg of Kdo_2_-Lipid A). The solution was then placed on ice and 10 mL cold acetone was added for every 1 mL hydrazine to precipitate ODLA followed by centrifugation for 30 min at 11,000 rpm. The pellet was washed 3 times each with cold acetone and diethyl ether; solvent was carefully decanted. After allowing the diethyl ether to evaporate ~5 min on its side under the fume hood, the pellets were dissolved in water and pooled together, flash frozen and lyophilized. Deacylation was confirmed by ESI-mass spectroscopy in 50:50 acetonitrile:water by observation of the half mass (m/z^−2^= 695).

### HepI Expression and Purification

HepI cloned from the *E. coli* K12 strain MB1760 (Escherichia coli ATCC 19215) was expressed in *E. coli* One Shot BL-21-AI as described previously with some changes. Two liters of LB-Amp media was inoculated with 10 mL of an overnight culture and allowed to grow at 37 °C, 200 rpm until an OD_600_ of 0.4-0.8 was reached (approximately 3-5 hrs). The cells were induced to a final concentration of 1 mM IPTG and 0.002% L-arabinose at 30 °C and expressed for 24 h. Cells were harvested by centrifugation for 10 min at 5,000 rpm. Supernatant was discarded and cell pellets were re-dissolved in 20 mL of binding buffer (20 mm HEPES, 1 mm imidazole and 500 mm NaCl, pH 7.4) for every liter of grown culture, to which ~1 mg of lysozyme was also added to aid in the lysing of cells. Cells were incubated on ice while mixing for 30 min, followed by homogenization at 18,000 psi for 5-10 cycles. The lysate was clarified by centrifugation at 13,000 rpm for 1 hr and the supernatant was loaded on to a Toyopearl AF-chelate-640 column attached to an ÄKTA purifier stored at 4 °C. The purification method charges the column with 500 μM cobalt sulfate, equilibrates with binding buffer, then protein is loaded and washed again with binding buffer. To remove any unbound protein the column was washed with wash buffer (20 mM HEPES, 40 mM imidazole and 500 mM NaCl, pH 7.4). Finally, HepI is eluted with strip buffer (50 mM ethylenediaminetetraacetic acid (EDTA) and 500 mL NaCl, pH 6.8).

To determine which samples contained HepI, SDS-PAGE gels stained with Coomassie blue stain were used to determine which fractions contain the 37 kDa protein (HepI). HepI consistently is co-eluted in the strip buffer with cobalt; fractions were pooled together and concentrated using 10,000 MWCO Vivaspin Ultra centrifugal concentrator. Concentrated (~ 5-10 mL) HepI was then placed over Bio-Scale Mini Bio-Gel P-6 desalting cartridge to buffer exchange into HepI storage buffer (100 mM HEPES, 1 M KCl, pH 7.5), column was also attached to an Äkta purifier at 4 °C. Again, fractions that contained purified protein were combined and concentrated using 10,000 MWCO Vivaspin Ultra centrifugal concentrator.

Protein was then stored in an amber, glass vial as a 50% ammonium sulfate precipitate at 4 °C. Protein remains stable for a few months after purification. To use for desired assay, precipitate was centrifuged at 4 °C for 6 min at 13,000 rpm, supernatant was removed and pellet was dissolved in desired buffer depending on experiment(s) to be carried out. Concentration was determined via nanodrop using the absorbance at 280 nm, and Beer’s law was employed to determine concentration given the extinction coefficient of HepI 55,928 M^−1^cm^−1^.

### Enzymatic assays

As previously reported, an ADP/NADH coupled assay was used to monitor HepI activity by monitoring the absorbance change at 340 nm at 37 °C on a Cary Bio 100 UV-Vis Spectrometer. Under normal conditions, the assay buffer was composed of 50 mM HEPES, 50 mM KCl, 10 mM MgCl_2_, pH 7.5. The coupled enzyme reaction additionally contained 100 μM phosphoenolpyruvate, 100 μM NADH, 100 μM dithiothreitol (DTT) and 0.05 U/μL of both pyruvate kinase and lactate dehydrogenase. 100 μM ADPH was used when ODLA concentration was varied and 100 μM ODLA was used when ADPH was varied. Once a stable baseline was established (~5 min), the reaction was initiated by addition of HepI to a final concentration of 50 nM or 100 nM (for high and lower substrate concentrations respectively) and all reported reaction rates are after background subtraction. When running 12 samples at once, plastic cuvette stirs and a repeat pipettor was used to inject 200 μL enzyme. Inhibition of HepI was monitored using the same ADP/NADH coupled assay, all conditions of the assay were the same as previously described unless otherwise stated. *K*_*i*_ values for each of the compounds were determined by varying compound final concentration from 100 μm to 10 nm, and a final concentration of 10 μM ODLA. Competition assays against ADPH were done with excess ODLA (100 μM) and varying ADPH from 2 - 0.25 μM. Competition assays against ODLA were done with excess ADPH (100 μM) and varying ODLA from 2 - 0.25 μM. Reactions were started with addition of enzyme and inhibitor was varied from 30 - 0.01 μM. Kinetics were fit to Lineweaver-Burke (Equation 1), competitive (Equation 2), non-competitive (Equation 3), mixed (Equation 4), Dixon (Equation 5), or Cornish-Bowden (Equation 6) models. All fitting and plotting was done through MATLAB R2018A.

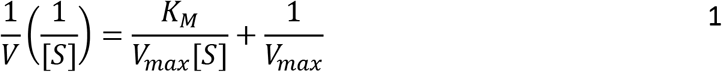

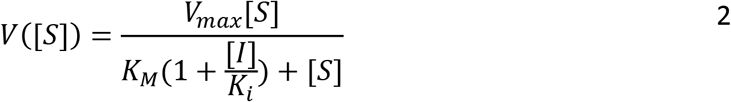

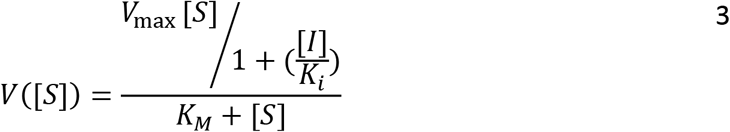

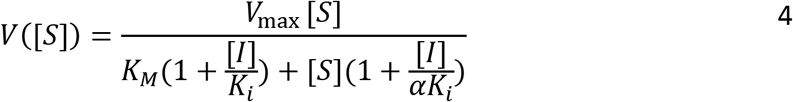

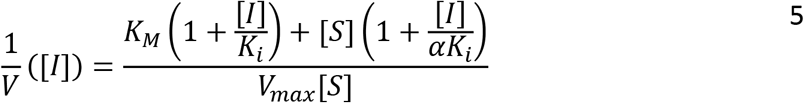

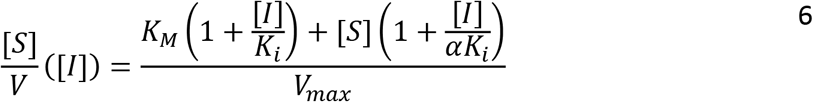

### Circular Dichroism Experiments

Circular dichroism melt experiments were performed in a Jasco J-810 spectropolarimeter in Starna 21-Q-2 cells (minimum volume 400 μL) sealed with Teflon tape to prevent evaporation. Parameters include 3 accumulations of data collected between 195-275 nm as a function of temperature (10-95 °C with 5 ° increments) at a ramp rate of 2 °/min and a scanning rate of 50 nm/min. Samples were prepared by diluting a mixture of 5 μM HepI and some combination of 250 μM substrate/product and/or 100 μM inhibitor in a buffer of 10 mM Tris-HCl and 100 mM KCl (pH = 7.5). CD data was analyzed via in house script written in MATLAB R2018A that average a set of triplicate data, normalizes to a percent unfolded (assuming the protein is fully folded at 5 °C) at 222nm as a function of temperature and fits to a sigmoid curve to determine the melt temperature (T_M_).

### Intrinsic tryptophan fluorescence spectra measurements

Fluorescence spectra were measured in triplicate at room temperature using 200 μL samples containing 1 μm HepI, in a 10 mm HEPES, 50 mm KCl, and 10 mm MgCl_2_, pH=7.5 buffer. Samples were contained in Starna 45-Q-3 cells (minimum volume 200μL). Substrate and inhibitor concentrations were 100 μM. All measurements were taken using a Fluoromax-4 fluorometer with an excitation slit bandpass of 2 nm and an emission slit bandpass of 3 nm (λ_ex_ = 290 nm, λ_em_ =310-450 nm). The data was subtracted from blanks, normalized and fit to a lognormal distribution (Equation 5)(26) to determine the emission maximum (λ_max_) with in house scripts written in MATLAB R2018A.

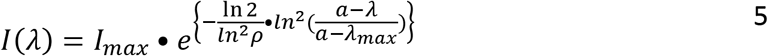

### Docking

HepI model system for docking and molecular dynamic simulations were prepared as previously described. Briefly, the HepI•ADPHep binary complex was modeled with the previously solved structure (PDB: 2H1H).(11) The ADP-2-deoxy-2-fluoro heptose present in this structure was modified to mimic the native donor via replacement of the 2-fluoro group with a hydroxyl group and inversion of the stereo configuration. The ternary complex was modeled with the previously solved pseudo-ternary complex (PDB: 6DFE).(27) This structure contains a native acceptor analogue with acyl chains replaced by singular acyl groups at the N positions of the terminal glucosamines. Furthermore, the acceptor is a phosphonate derivate and was replaced with the modified native donor from the HepI•ADPHep binary complex described above.

Structures were prepared in Maestro with the protein preparation wizard and provided default settings.(28) Missing loops and sidechains were modeled with Prime and protonation states were determined with PROPKA.(29–32) Docking was carried out in Glide with standard precision and the OPLS3e forcefield.(33,34) All resulting poses were subject to MMGBSA end point free energy approximation and the lowest energy pose was selected for molecular dynamic simulations.(35,36)

### Molecular Dynamic Simulations

Molecular dynamic simulations were implemented in the GROMACS 2021.1 (37) package with the AMBER99SB forcefield.(38) Ligands were parametrized with antechamber from AmberTools20.(39) Atom types were assigned from the second generation general AMBER forcefield (GAFF2) and charges were applied with the AM1-BCC method.(40,41) Resulting files were converted to Gromacs compatible file formats with the ACPYPE script.(42) The protein ligand complexes were placed into a dodecahedron with periodic boundary condition solvated with the TIP3P water model.(43) The system was electroneutralized with the addition of counterions and subsequent addition of ions to a concentration of 0.150 M to mimic physiological conditions. The steepest decent algorithm was used for energy minimization. The system was further equilibrated with subsequent 1 ns simulations under isochoric/isothermal (NVT) and isobaric/isothermal (NPT) conditions. During equilibration harmonic restraints (1000 kJ/mol/nm^2^) were applied to all heavy atoms and gradually removed in a stepwise fashion, initially from sidechains then the backbone, over the course of 10 ns. Production simulations were carried out for 100 ns in triplicate under isobaric/isothermal (NPT) conditions with a 2 fs timestep at 300 K and 1 atm. Long range electrostatic interactions were calculated with the particle-mesh-Ewald with a fourth order cubic interpolation and a 1.6 Å grid spacing. Short range nonbonded interactions were calculated with a 10 Å cutoff. Temperature was maintained with the V-rescale thermostat, while pressure was maintained with the Berendsen and Parrinello-Rahman barostat during equilibration or production simulations, respectively.(44–47) Bonds were constrained with the LINCS method.(48) Root mean square deviation (RMSD) of the protein backbone and root mean square fluctuations (CαRMSF) of the Cα were calculated in GROMACS for individual trajectories. Averages and standard deviations of the RMSD and CαRMSF were calculated and plotted via Python3.(49)

### Binding Free Energies

Binding energies of inhibitors to the protein/substrate ternary complex were determined by the molecular mechanics poisson boltzman solvent accessible (MMPBSA) method in Amber using the MMPBSA.py script and gmx_MMPBSA extension with an internal dielectric constant of 4. (50,51) The binding energies and its standard deviation were evaluated from 300 representative frames taken at 100ps time interval for each of the three 100ns simulated trajectories. Experimental binding energies of the inhibitors to the protein/substrate complex were estimated with the Gibbs Free Energy equation and the equilibrium constant approximated as Ki. Both values were plotted as a function of another and fit to straight line to determine a correlation value.

## Abbreviations

GT: glycosyltransferase
CAZY: Carbohydrate-Active enZYmes Database
HepI: Heptosyltransferase I
ADP-Hep: ADP-L-*glycero*-β-D-*manno*-heptose
Kdo: 3-deoxy-D-*manno*-oct-2-ulosonic acid
ODLA: O-deacylated Kdo_2_-Lipid A
H-Kdo_2_-Lipid A: heptosylated Kdo_2_-Lipid A
FDLA: fully-deacylated Kdo_2_-Lipid A
ADP: adenosine disphosphate
FDHLA: fully-deacylated heptosylated-Kdo_2_-Lipid A
FDLA-H: deprotonated sugar donor nucleophile
D13+H: protonated aspartic acid 13
TIP3P: transferrable intermolecular potential with 3 points
RMSD: root mean square deviation
Rgyr: radius of gyration
RMSF: root mean square fluctuations
PCA: Principal component analysis
DCCM: dynamic cross correlation matrix
Amp: Ampicillin
Tet: Tetracycline
CD: Circular dichroism
Tm: melting temperature

## Acknowledgements

We would like to acknowledge the University of Minnesota Supercomputing Institute provided the schrodinger software package for performing the docking studies.

## Data availability statement

*The data underlying this article are available in the article and in its online supplementary material. Software used in this study are available from their respective sources. Scripts and simulation trajectories are available from the corresponding authors upon request.*

