## Supplemental Information for "Discovery of Aminoglycosides as first in class, nanomolar inhibitors of Heptosyltransferase I" (from another version of this preprint: version 1, 2021-10-04)

Inhibiting HepI

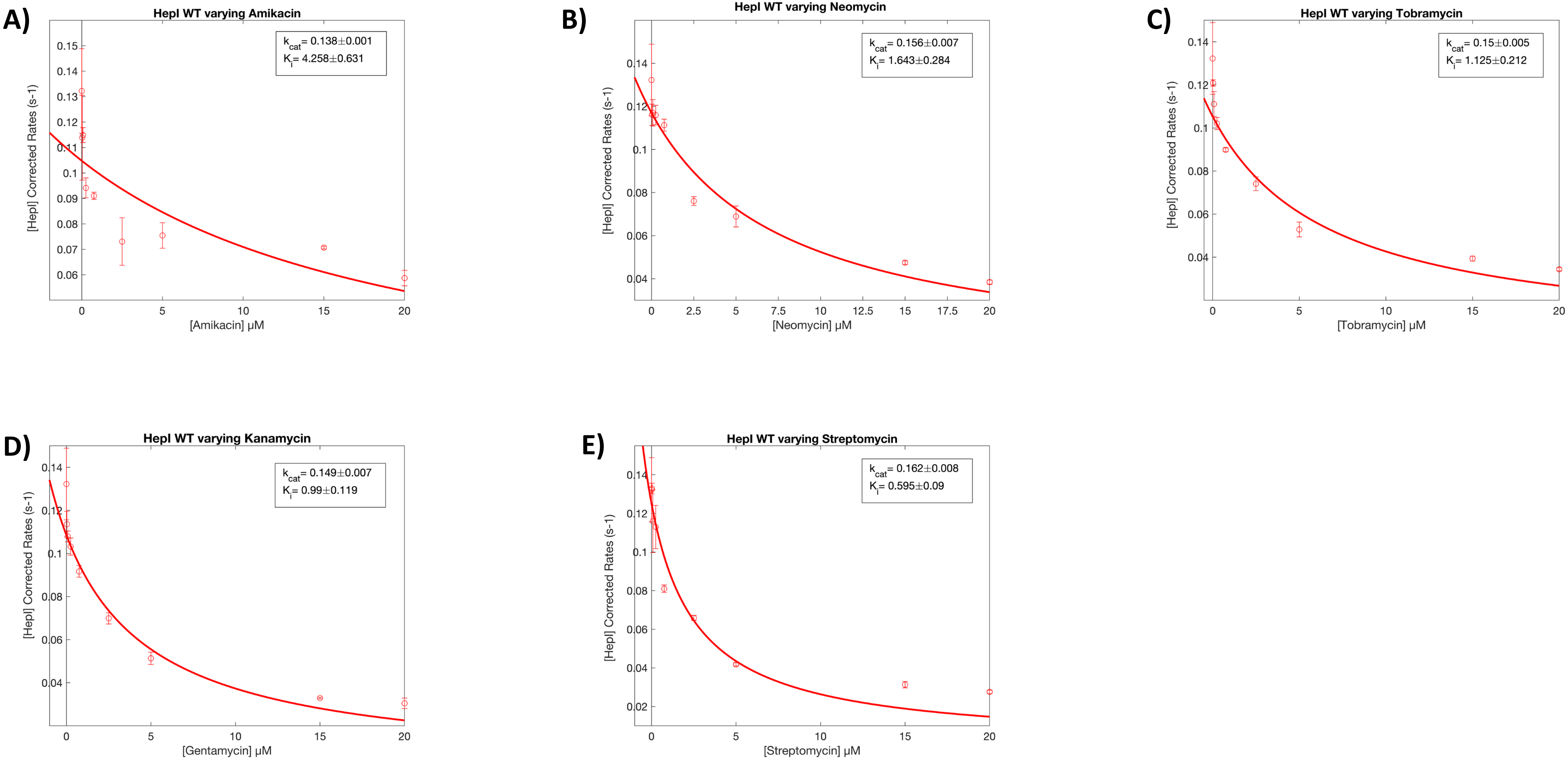


Supplemental Figure 1: Inhibition kinetic curves for HepI in the presence of **A)** Amikacin, **B)** Neomycin, **C)** Tobramycin, **D)** Kanamycin B, or **E)** Streptomycin.

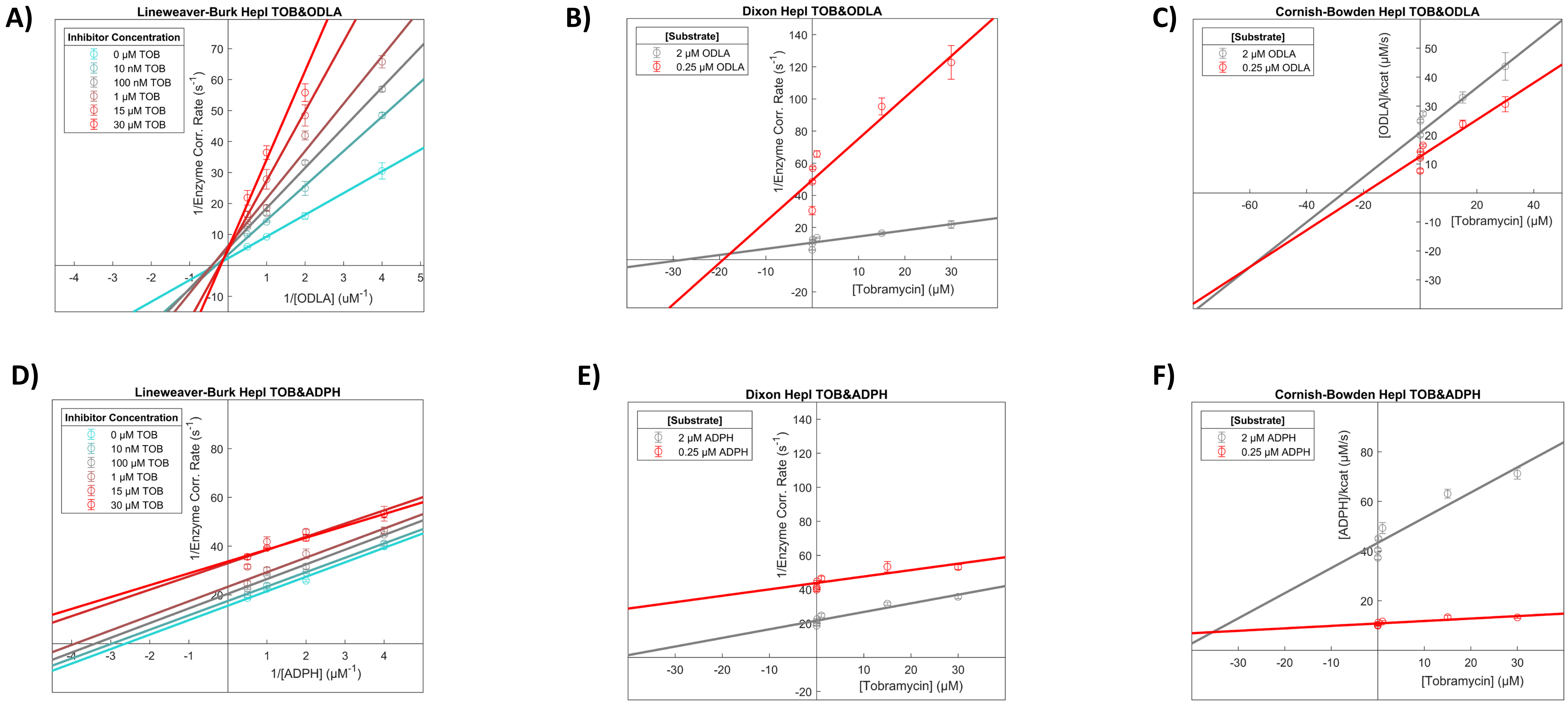


Supplemental Figure 2: Double reciprocal plots of HepI kinetics in the presence of tobramycin with varying **A-C)** ODLA or **D-F)** ADPH.

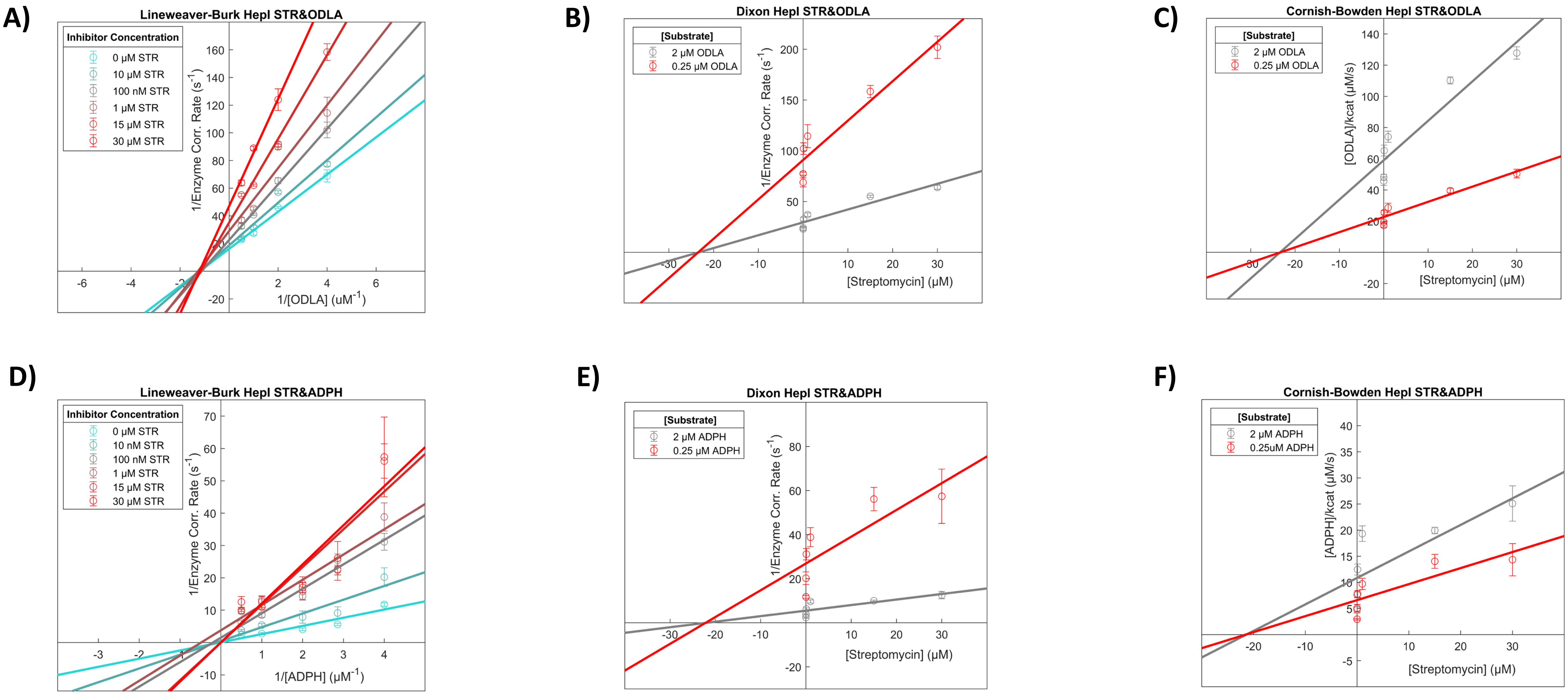


Supplemental Figure 3: Double reciprocal plots of HepI kinetics in the presence of streptomycin with varying **A-C)** ODLA or **D-F)** ADPH.

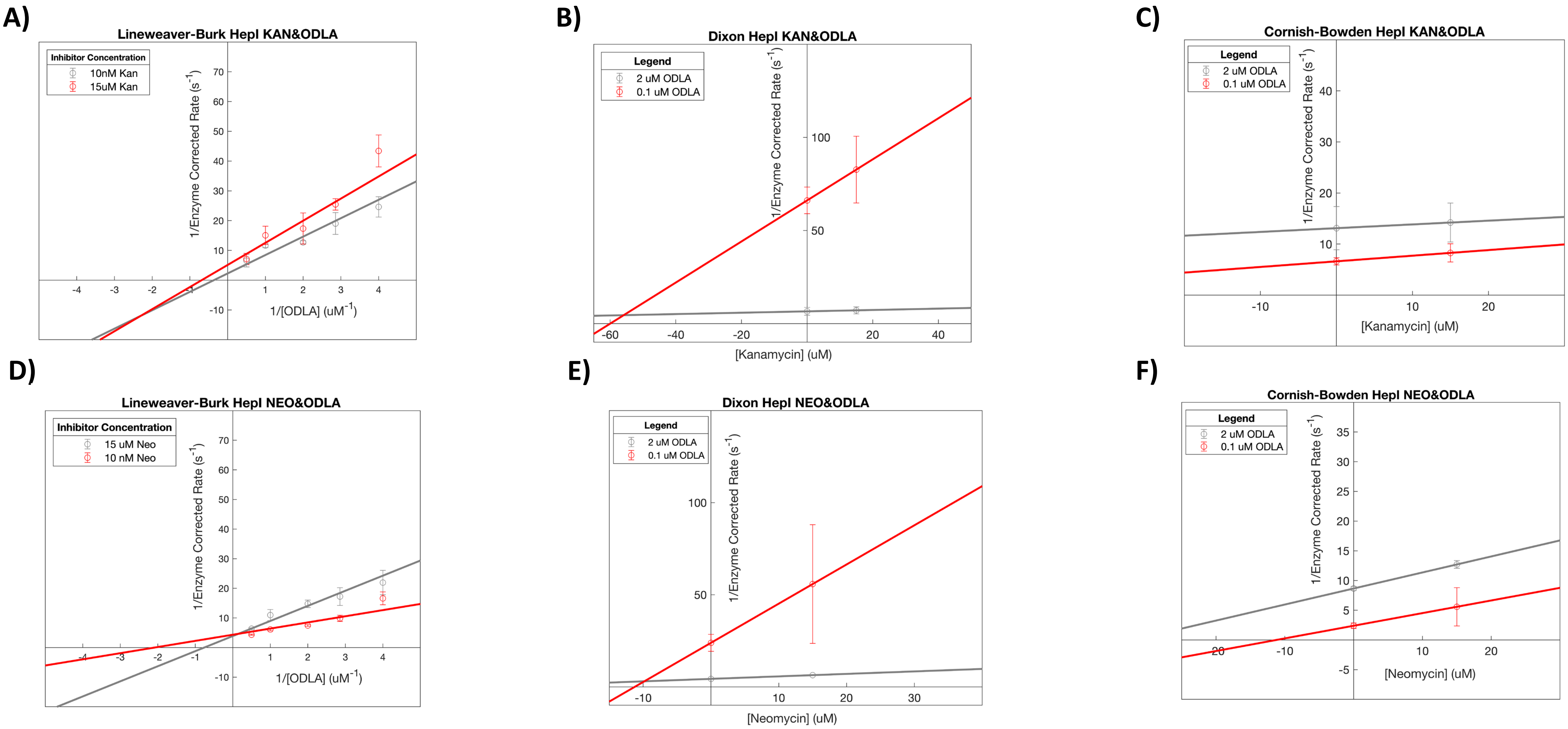


Supplemental Figure 4: Double reciprocal plots of HepI kinetics in the presence of ODLA with varying **A-C)** Kanamycin B or **D-F)** Neomycin.

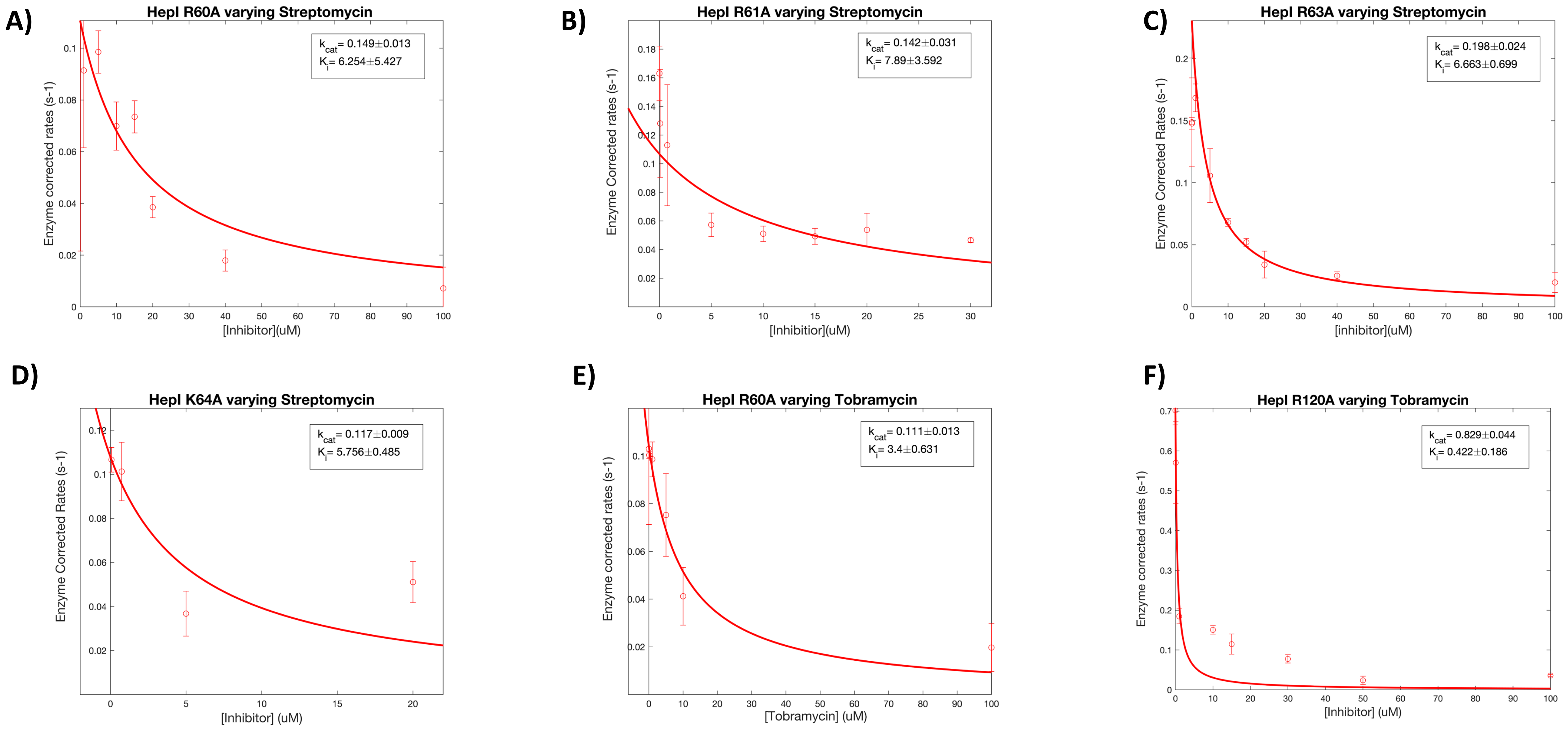


Supplemental Figure 5: Inhibition kinetic curves for HepI mutants **A)** R60A, **B)** R61A, **C)** R63A, **D)** and K64A in the presence of streptomycin. HepI inhibition kinetic curves for **E)** R60A, and **F)** R120A in the presence of tobramycin.

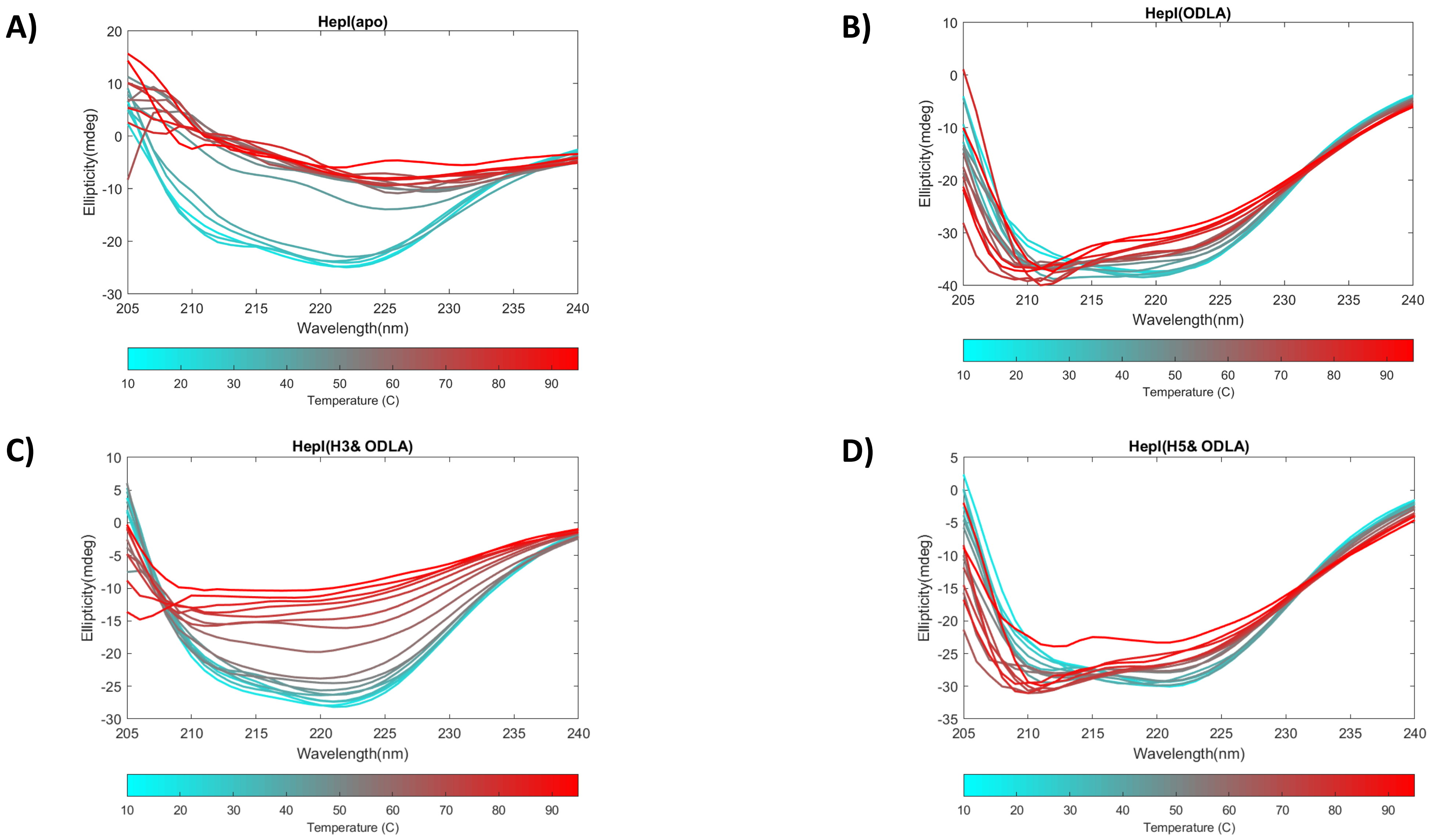


Supplemental Figure 6: CD melts of HepI **A)** apo or in the presence of **B)** ODLA, **C)** ODLA/tobramycin, and **D)** ODLA/streptomycin.

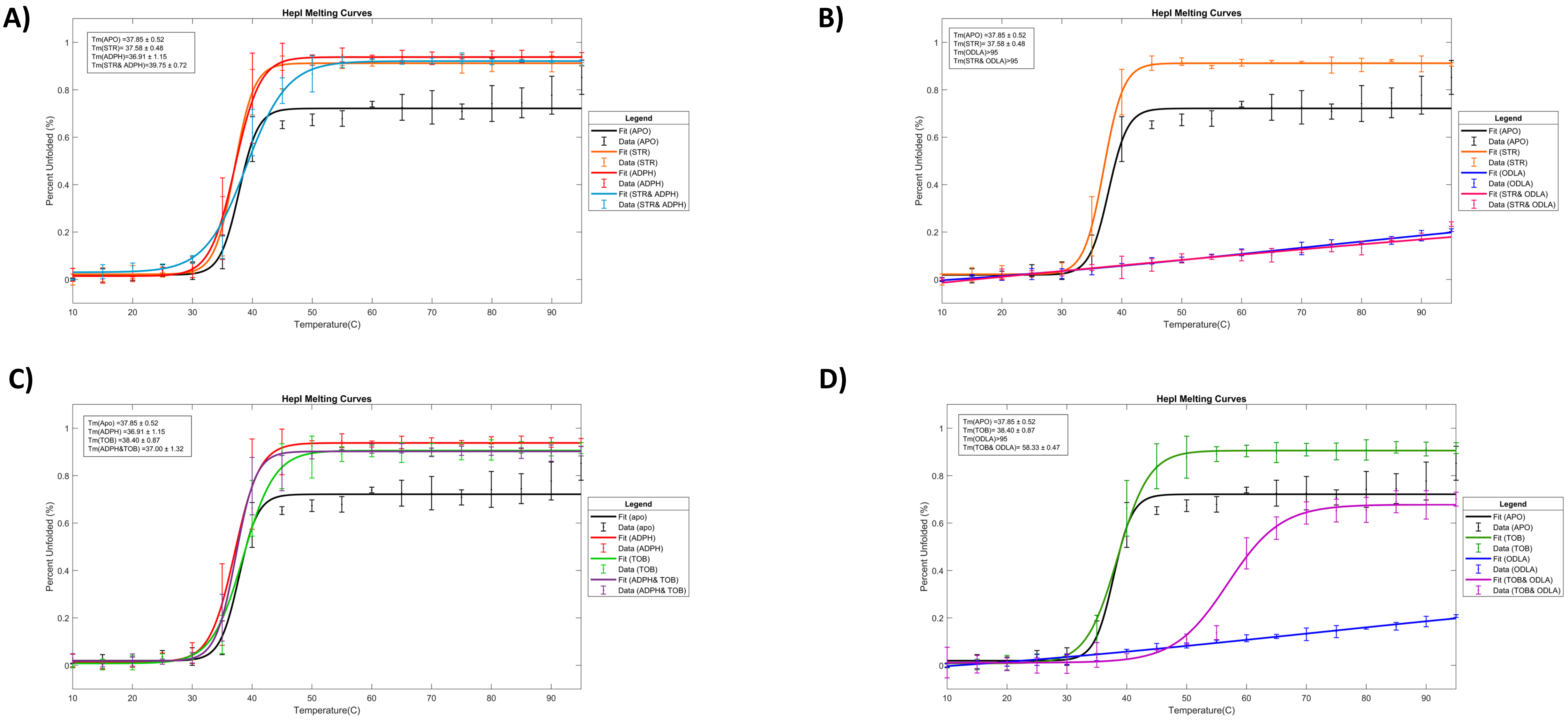


Supplemental Figure 7: Wildtype HepI unfolding monitored via CD at 222nm across several temperatures for **A)** streptomycin with ADPH, **B)** streptomycin with ODLA, **C)** tobramycin with ADPH and **D)** tobramycin with ODLA.

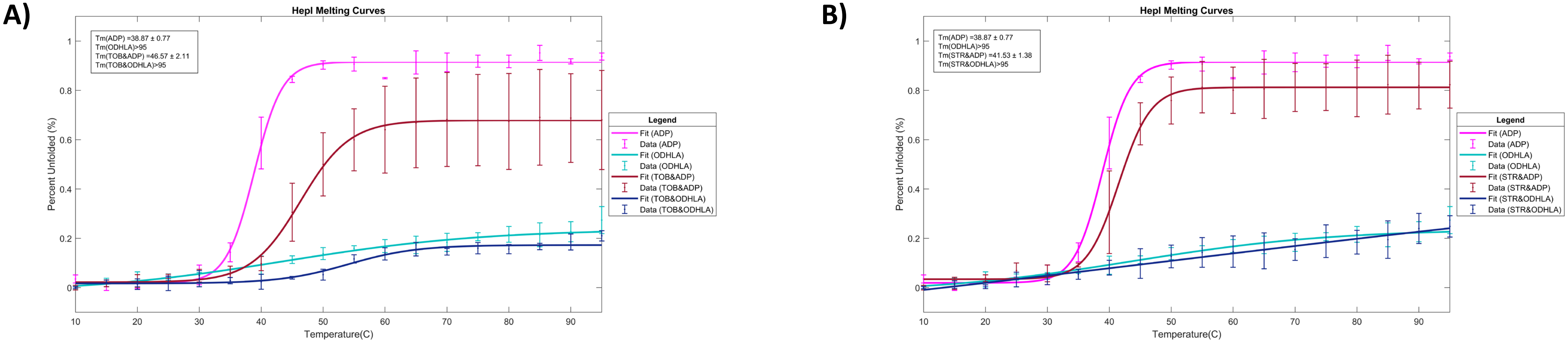


Supplemental Figure 8: HepI unfolding monitored via CD at 222nm across several temperatures for **A)** tobramycin with products and **B)** streptomycin with products.

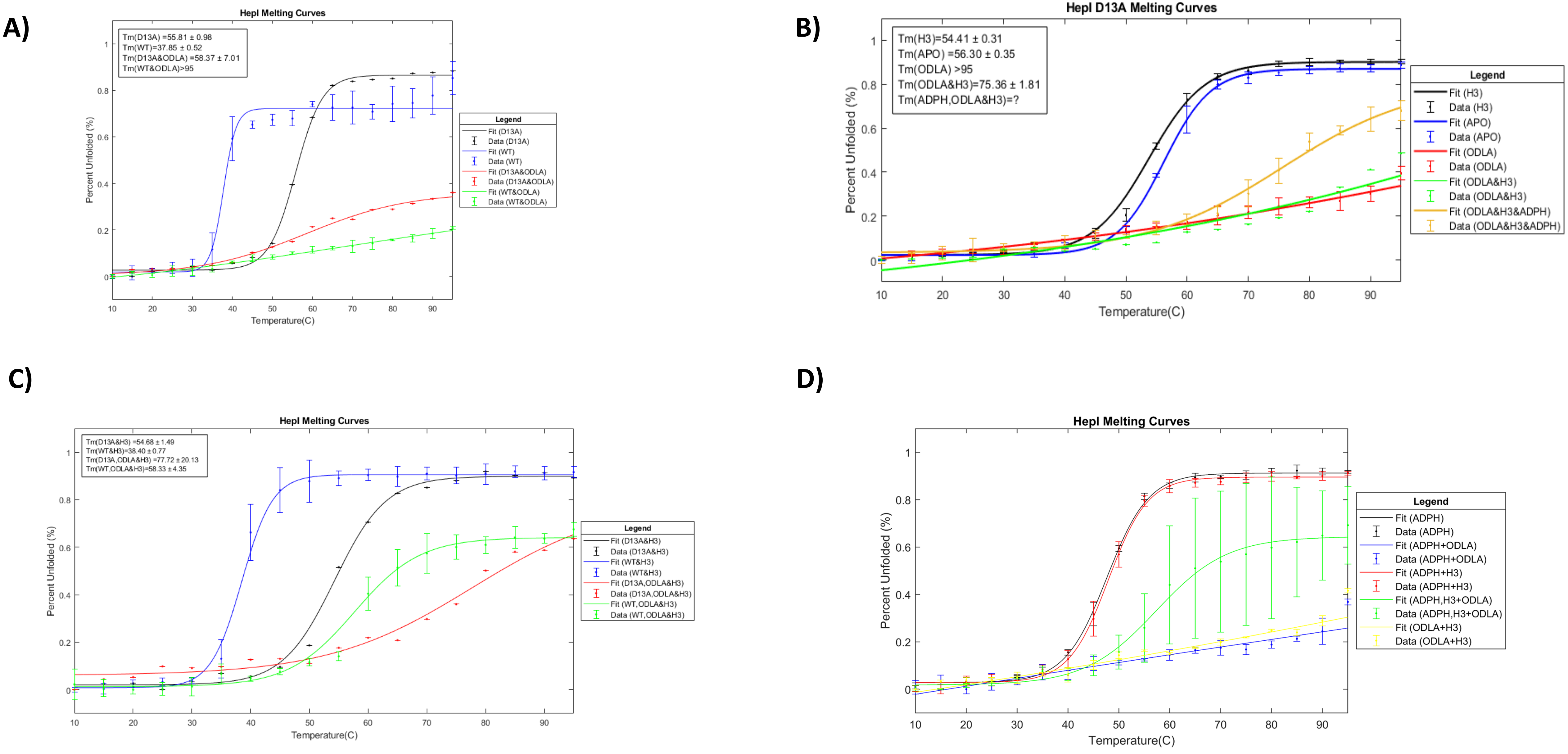


Supplemental Figure 9: HepI unfolding monitored via CD at 222nm across several temperatures for **A)** wildtype and D13A with substrates, **B)** D13A with substrates and tobramycin, **C)** D13A with substrates and tobramycin and **D)** D13A with substrates and tobramycin.

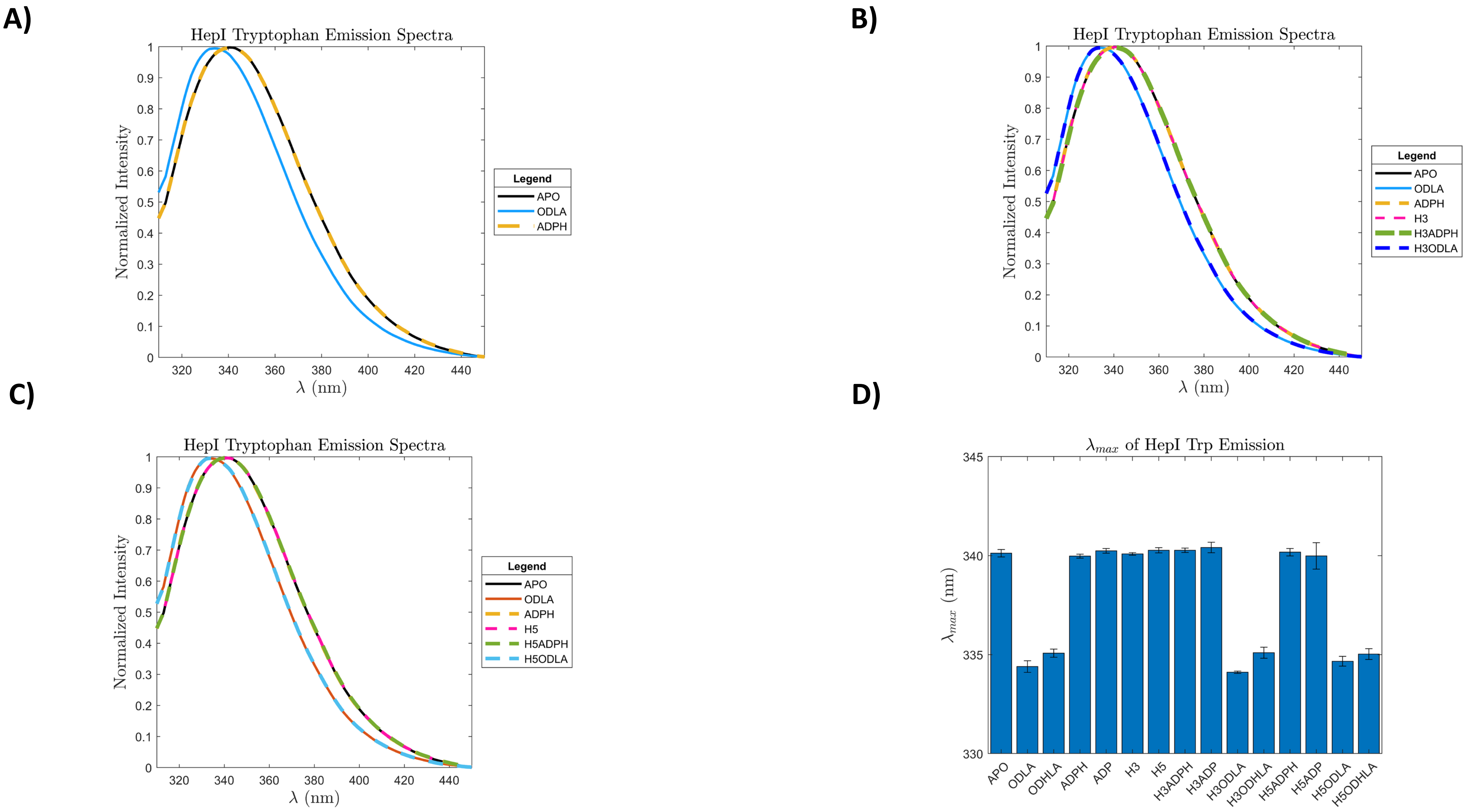


Supplemental Figure 10: HepI tryptophan fluorescence emission spectra for **A)** substrates, **B)** with tobramycin, **C)** with streptomycin, and **B)** $\lambda_{max}$ in the presence/absence of substrates, products and inhibitors.

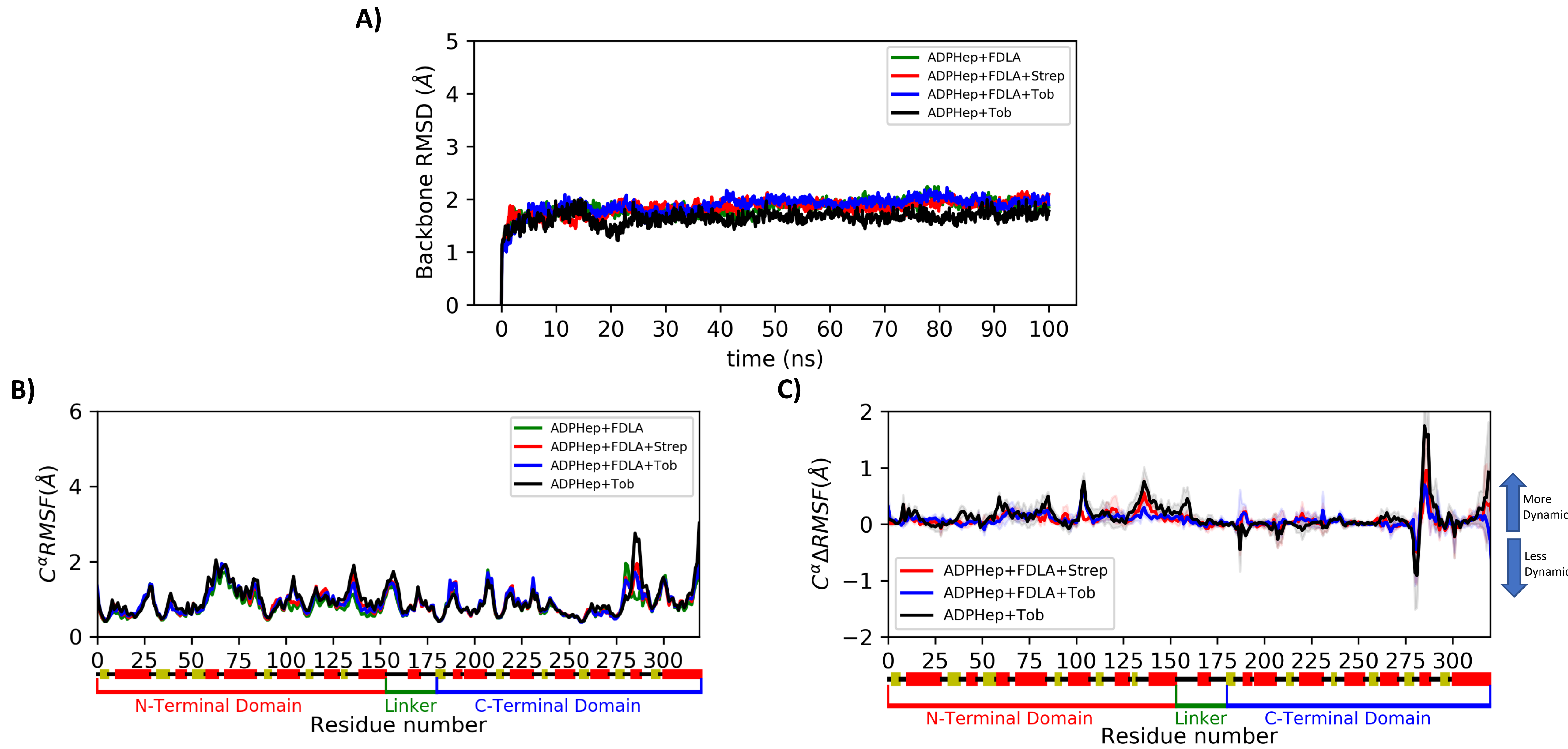


Supplemental Figure 11: (**A**) Backbone RMSD, (**B**) $C_{\alpha}$ RMSF and (**C**) $C_{\alpha}$ ΔRMSF of HepI substrate complex (green), in the presence of Streptomycin (red) or Tobramycin (blue) and HepI binary complex in the presence of Tobramycin(black). Secondary structure of residues are indicated by color (red:α-helix, yellow:β-sheet, black:random coil) below their respective positions.

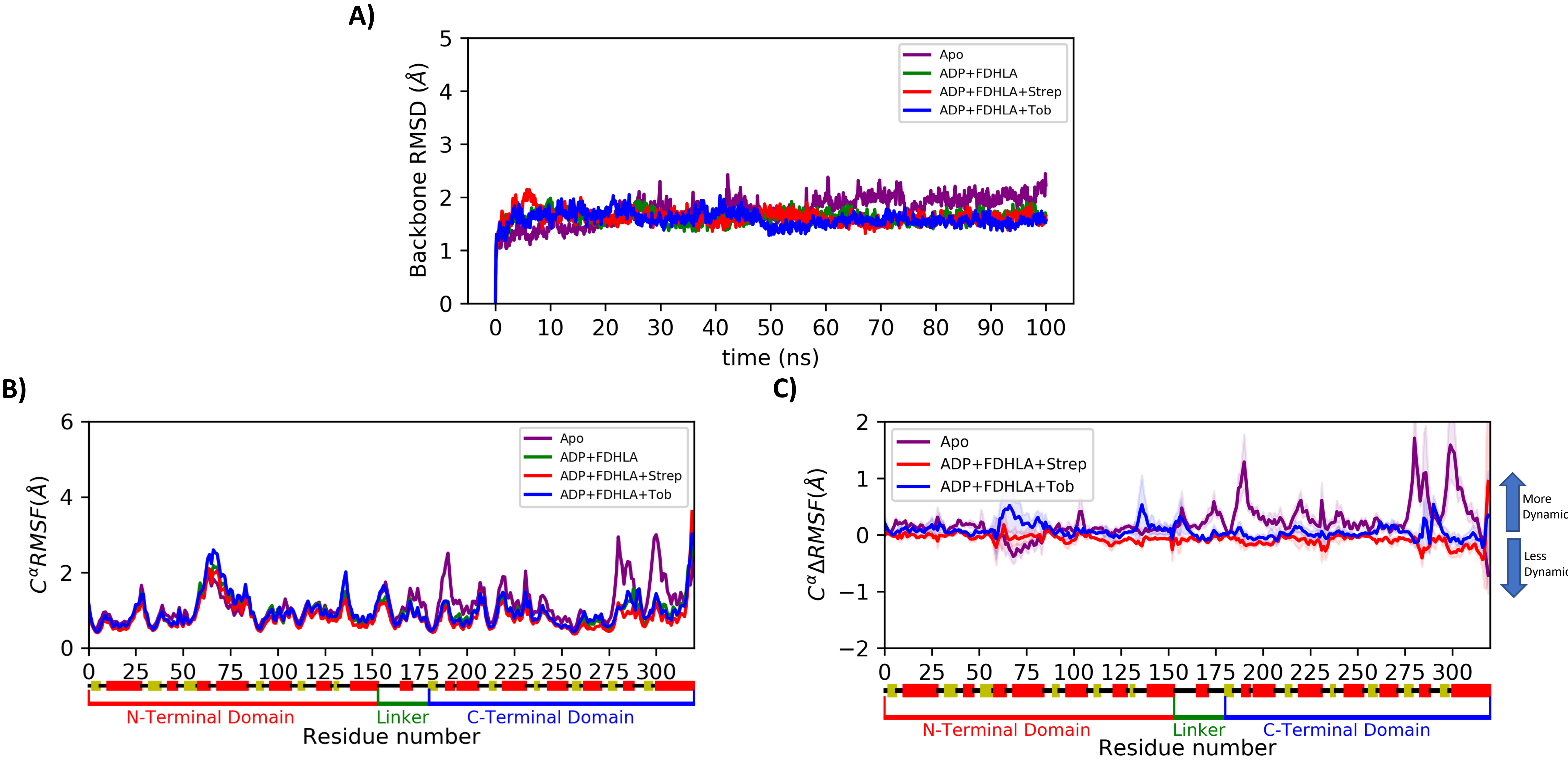


Supplemental Figure 12: (**A**) Backbone RMSD, (**B**) $C_{\alpha}$ RMSF and (**C**) $C_{\alpha}$ ΔRMSF of HepI apo (purple), substrates (green), in the presence of Streptomycin (red) or Tobramycin (blue). Secondary structure of residues are indicated by color (red:α-helix, yellow:β-sheet, black:random coil) below their respective positions.

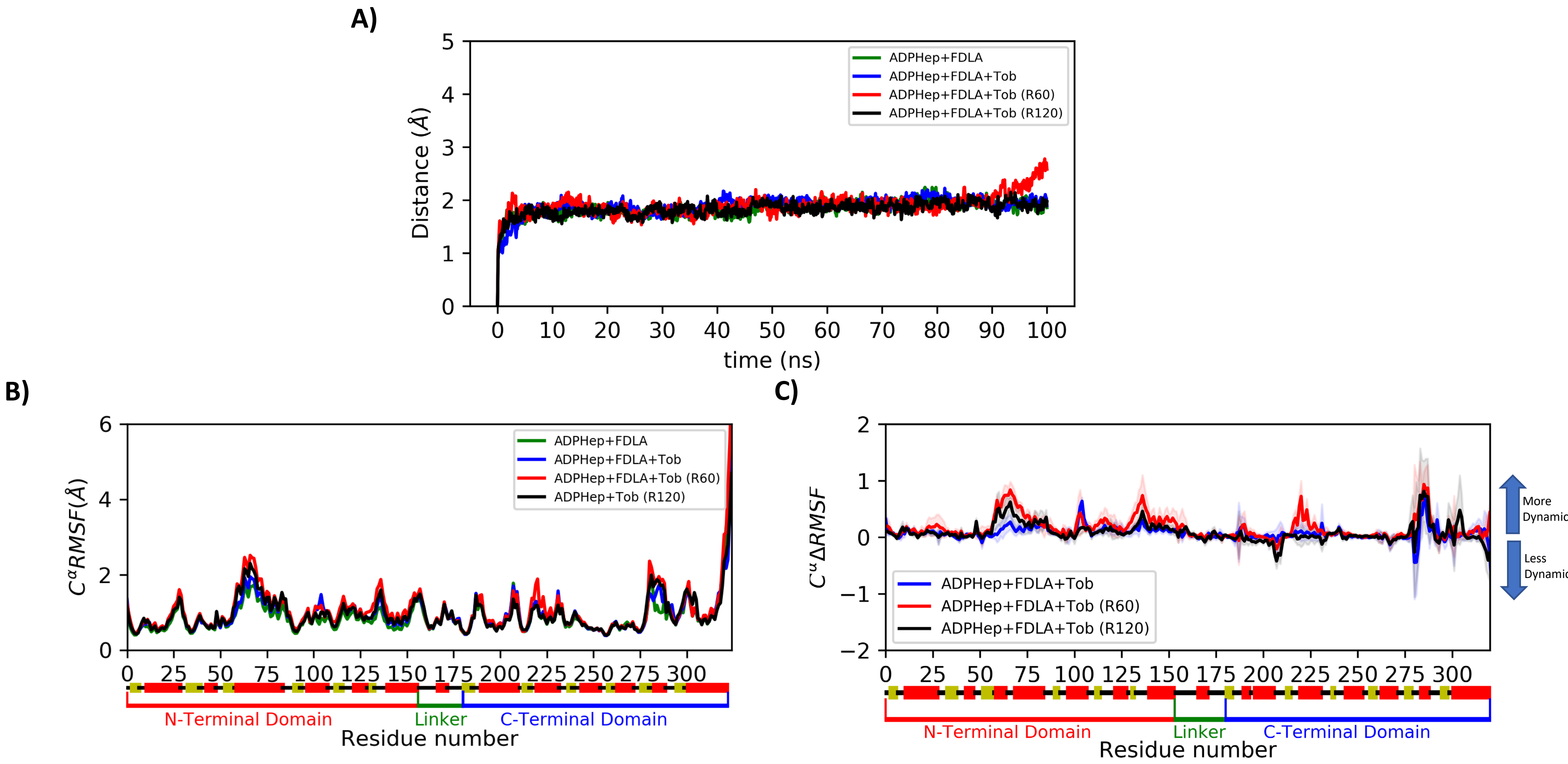


Supplemental Figure 13: (**A**) Backbone RMSD, (**B**) $C_{\alpha}$ RMSF and (**C**) $C_{\alpha}$ ΔRMSF of HepI (WT) with substrates (green), in the presence of Tobramycin (blue), HepI (Arg60A) with substrates in the presence of Tobramycin (red), HepI (Arg120A) with substrates in the presence of Tobramycin (black) . Secondary structure of residues are indicated by color (red:α-helix, yellow:β-sheet, black:random coil) below their respective positions.

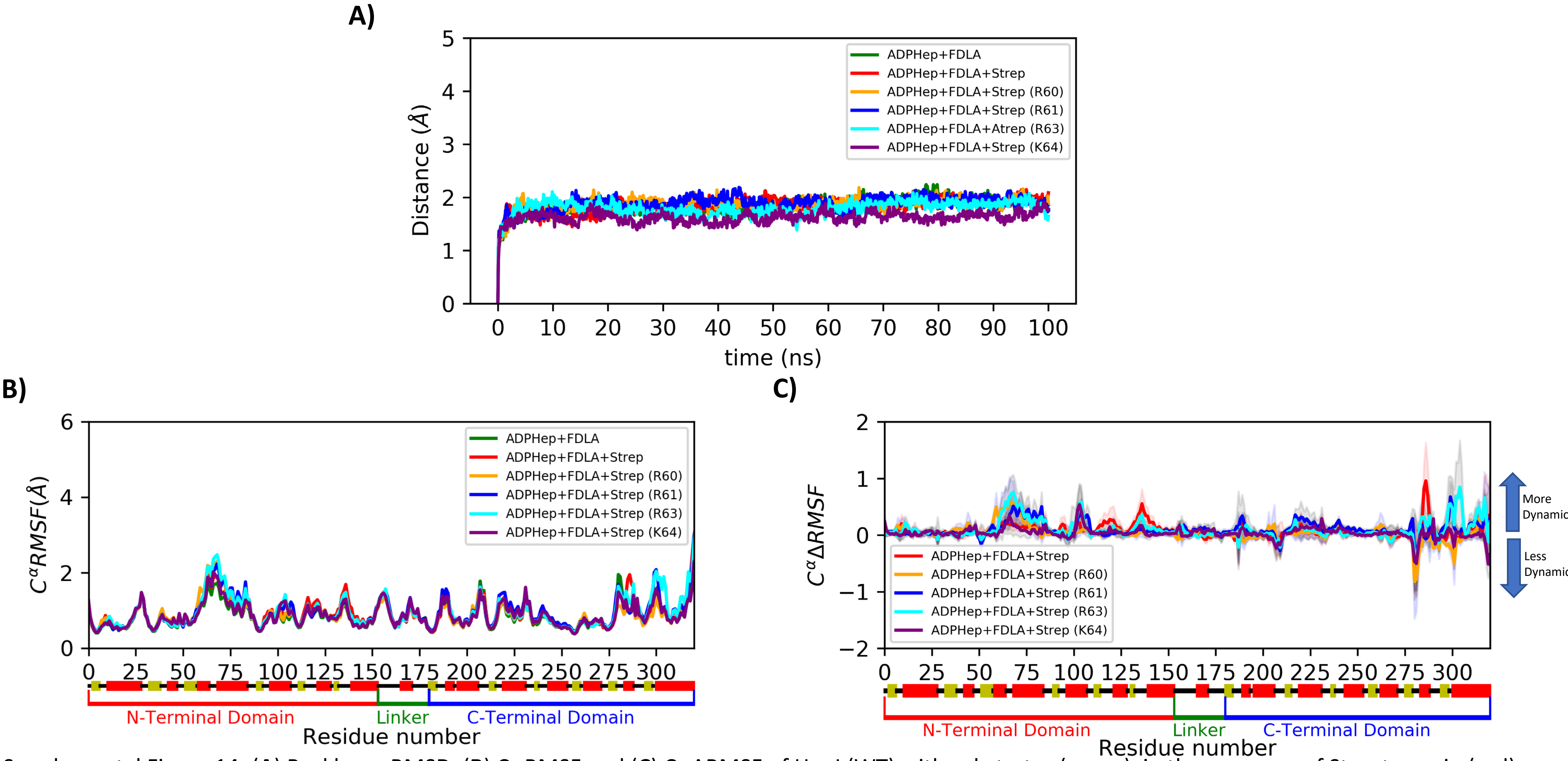


Supplemental Figure 14: (**A**) Backbone RMSD, (**B**) $C_{\alpha}$ RMSF and (**C**) $C_{\alpha}$ ΔRMSF of HepI (WT) with substrates (green), in the presence of Streptomycin (red), HepI (Arg60A) with substrates in the presence of Streptomycin (orange), HepI (Arg61A) with substrates in the presence of Streptomycin (blue), HepI (Arg63A) with substrates in the presence of Streptomycin (cyan), and HepI (Lys64A) with substrates in the presence of Streptomycin (purple) . Secondary structure of residues are indicated by color (red:α-helix, yellow:β-sheet, black:random coil) below their respective positions.

A)

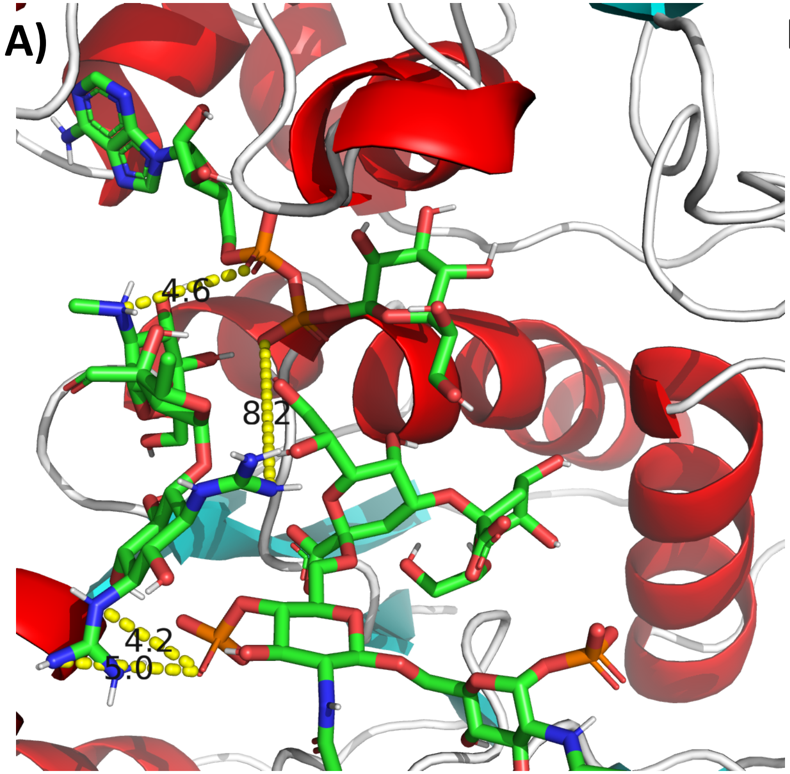


B)

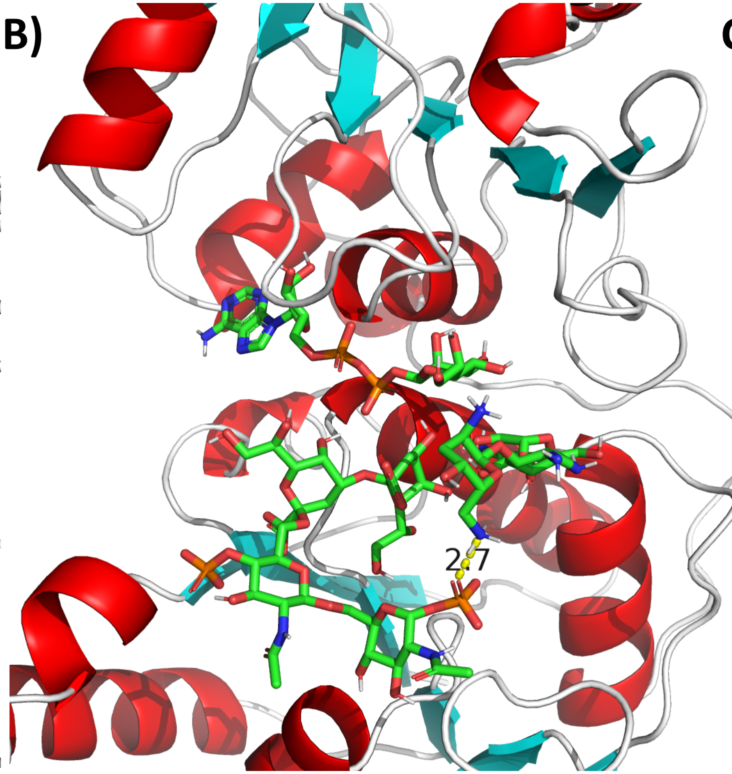


C)

| ADPHep | FDLA | Streptomycin | Tobramycin | Average Distance (Å) |
|---|---|---|---|---|
| Pα (-PO-) | | C2’’(-NH2+-) | | 3.7 ± 1.0 |
| Pβ(-PO-) | | C8’(-NH2) | | 7.6 ± 0.9 |
| | C4 Glcnac(-PO-) | C7’(=NH+-) | | 6.8 ± 1.1 |
| | C4 Glcnac (-PO-) | C7’(-NH2) | | 6.8 ± 1.5 |
| | C1 Glcnac (-PO-) | | C6’’ (-NH3+) | 4.5 ± 0.9 |

Supplemental Figure 15: (**A**) Model of interactions between ADPHep/FDLA with Streptomycin and (**B**) model of interactions between ADPHep/FDLA with Tobramycin and **C**) average distance of these interactions across three independent trajectories.

A)

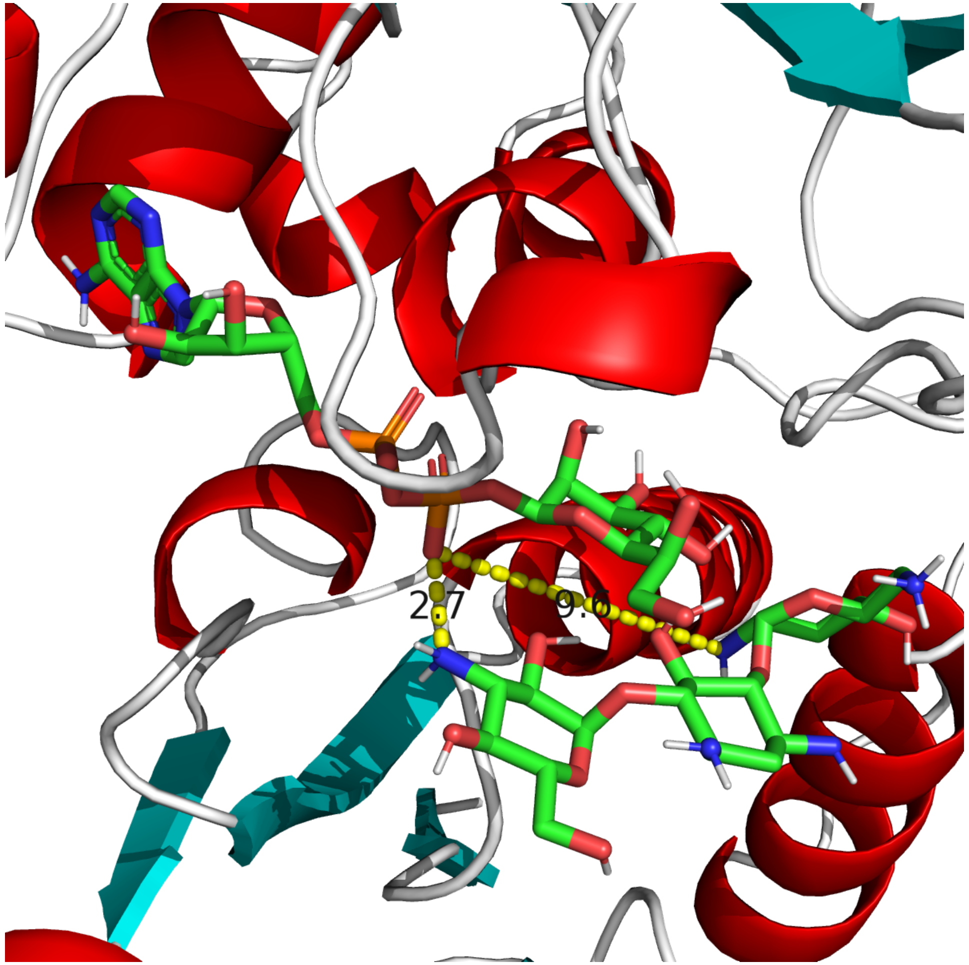


B)

| ADPHep | Tobramycin | Average Distance (Å) |
|---|---|---|
| O1B (-PO-) | N4 (-NH3+) | 3.3 ± 0.3 |
| O1B (-PO-) | N3(-NH3+) | 8.8 ± 0.4 |
| O3(-OH) | O4’(-OH) | 2.8 ± 0.1 |

Supplemental Figure 16: (**A**) Representative model and **(B)** table of interactions between ADPHep with Tobramycin  across three independent trajectories.

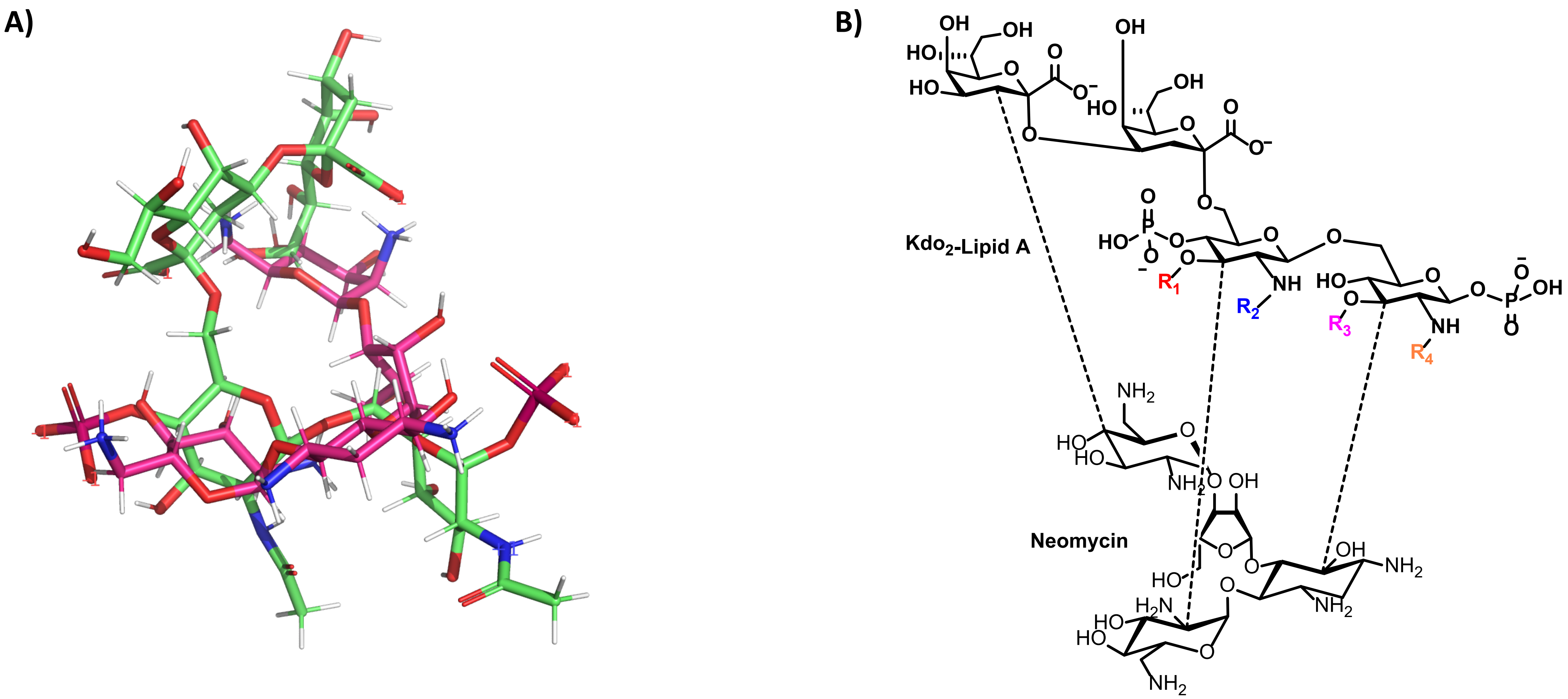


Supplemental Figure 17: (**A**) 3D and **(B)** 2D Structural superposition of Neomycin on fully deacylated donor substrate that neomycin acts as competitive inhibitor towards. Dotted lines show which sugars align with another.

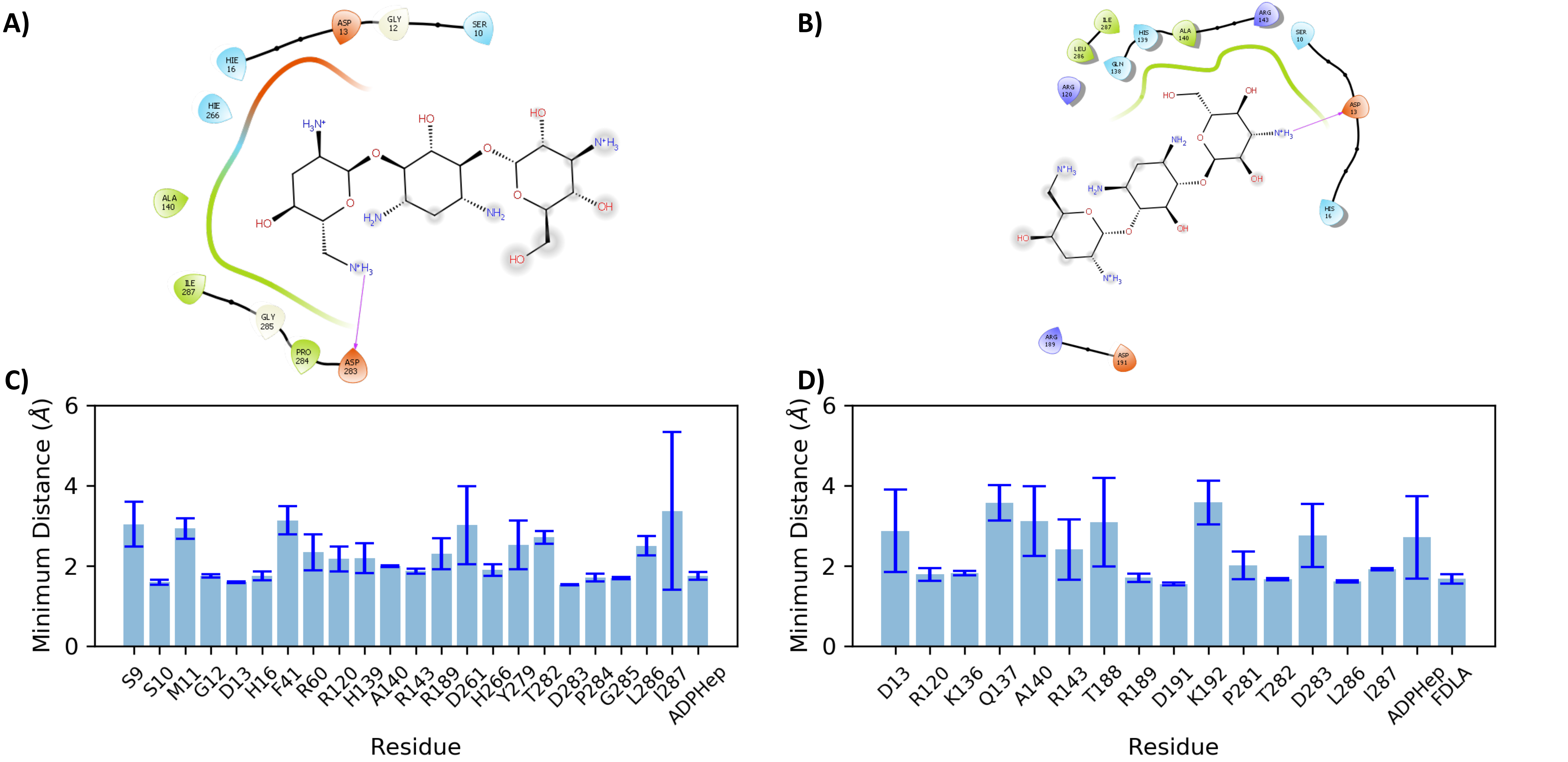


Supplemental Figure 18: (**A**) Tobramycin (binary) Ligand Interaction Diagram (**B**) Tobramycin (ternary) Ligand Interaction Diagram (**C**) Residues with minimum distance <3.5 Å from Tobramycin (binary) and (**D**) Residues with minimum distance <3.5 Å from Tobramycin (ternary)

A)

| Simulation | PDB Code | Mutation | ADP-Hep | FDLA | ADP | FDHLA | Tob | Strep |
|---|---|---|---|---|---|---|---|---|
| HepI | 2GT1 | WT | N/A | | | | | |
| ADP-Hep•FDLA•HepI | 6DFE | WT | Added | Present | N/A | | | |
| ADP•FDHLA•HepI | 6DFE | WT | N/A | | Added | Modified | N/A | |
| Tob•ADP-Hep•FDLA•HepI | 6DFE | WT | Added | Present | N/A | | Docked | N/A |
| Strep•ADP-Hep•FDLA•HepI | 6DFE | WT | Added | Present | N/A | | N/A | Docked |
| Tob•ADP-Hep•HepI | 2H1H | WT | Modified | N/A | N/A | | Docked | N/A |
| Tob•ADP•FDHLA•HepI | 6DFE | WT | N/A | | Added | Modified | Docked | N/A |
| Strep•ADP•FDHLA•HepI | 6DFE | WT | N/A | | Added | Modified | N/A | Docked |
| Tob•ADP-Hep•FDLA•HepI | 6DFE | Arg60Ala | Added | Present | N/A | | Docked | N/A |
| Tob•ADP-Hep•FDLA•HepI | 6DFE | Arg120Ala | Added | Present | N/A | | Docked | N/A |
| Strep•ADP-Hep•FDLA•HepI | 6DFE | Arg60Ala | Added | Present | N/A | | N/A | Docked |
| Strep•ADP-Hep•FDLA•HepI | 6DFE | Arg61Ala | Added | Present | N/A | | N/A | Docked |
| Strep•ADP-Hep•FDLA•HepI | 6DFE | Arg63Ala | Added | Present | N/A | | N/A | Docked |
| Strep•ADP-Hep•FDLA•HepI | 6DFE | Lys64Ala | Added | Present | N/A | | N/A | Docked |

Supplemental Table 1: The simulations performed in this study including PDB codes of crystal structures that were used and whether ligands were present in the original structure, added into position based on information from another crystal structure, modified atoms to mimic the native ligands, docked or if any mutations were introduced into the structure.

| Simulation | Mutation | RMSD (Å) | $R_{gyr}$ (Å) | Interdomain Distance (Å) | RMSF (Residues >1.5 Å) |
|---|---|---|---|---|---|
| HepI | WT | 1.76 ± 0.45 | 21.32 ± 0.24 | 30.26 ± 0.67 | 29, 60, 63, 64, 65, 66, 67, 68, 69, 70, 157, 158, 171, 175, 188, 189, 190, 191, 192, 208, 210, 218, 219, 220, 221, 232, 280, 281, 282, 283, 284, 285, 286, 287, 288, 289, 298, 299, 300, 301, 302, 303, 304, 305, 306, 317, 318, 319, 320, 321, 322, 323 |
| ADP-Hep•FDLA•HepI | WT | 1.84 ± 0.23 | 21.13 ± 0.16 | 29.74 ± 0.43 | 64, 65, 67, 68, 69, 70, 208, 210, 281, 282, 283, 301, 302, 319, 320, 321, 322, 323, 324, 325 |
| ADP•FDHLA•HepI | WT | 1.62 ± 0.26 | 21.09 ± 0.16 | 28.24 ± 0.38 | 60, 61, 63, 64, 65, 66, 67, 68, 69, 70, 71, 72, 73, 84, 137, 289, 317, 318, 319, 320 |
| Tob•ADP-Hep•FDLA•HepI | WT | 1.88 ± 0.30 | 21.23 ± 0.12 | 30.05 ± 0.37 | 63, 64, 65, 66, 67, 68, 69, 71, 187, 207, 209, 231, 280, 284, 285, 286, 300, 301, 317, 318, 319, 320, 321, 322, 323, 324 |
| Strep•ADP-Hep•FDLA•HepI | WT | 1.84 ± 0.25 | 21.28 ± 0.16 | 30.28 ± 0.47 | 62, 63, 64, 65, 66, 67, 68, 69, 72, 135, 136, 207, 280, 285, 286, 287, 288, 300, 317, 318, 319, 320, 321, 322, 323, 324 |
| Tob•ADP-Hep•HepI | WT | 1.65 ± 0.30 | 21.26 ± 0.19 | 29.53 ± 0.56 | 59, 60, 62, 63, 64, 65, 66, 67, 68, 69, 72, 83, 84, 85, 104, 134, 135, 136, 137, 156, 157, 158, 207, 284, 285, 286, 287, 288, 300, 317, 318, 319 |
| Tob•ADP•FDHLA•HepI | WT | 1.59 ± 0.33 | 21.32 ± 0.9 | 28.76 ± 0.51 | 61, 62, 63, 64, 65, 66, 67, 68, 69, 70, 71, 72, 73, 74, 76, 77, 84, 136, 137, 156, 157, 158, 286, 291, 292, 318, 319, 320 |
| Strep•ADP•FDHLA•HepI | WT | 1.62 ± 0.30 | 21.16 ± 0.14 | 28.29 ± 0.34 | 63 , 64, 65, 66, 67, 68, 69, 70, 71, 72, 73, 318, 319 ,320 |
| Tob•ADP-Hep•FDLA•HepI | Arg60Ala | 1.90 ± 0.28 | 21.38 ± 0.18 | 30.45 ± 0.52 | 28, 59, 60, 61, 62, 63, 64, 65, 66, 67, 68, 69, 70, 71, 72, 73, 75, 76, 134, 135, 136, 137, 154, 157, 207, 219, 220, 280, 281, 282, 283, 284, 285, 286, 287, 288, 300, 301, 318, 319, 320, 321, 322, 323, 324 |
| Tob•ADP-Hep•FDLA•HepI | Arg120Ala | 1.83 ± 0.33 | 21.29 ± 0.19 | 30.19 ± 0.53 | 59, 60, 62, 63, 64, 65, 66, 67, 68, 69, 70, 71, 72, 83, 136, 280, 281, 282, 283, 284, 285, 286, 287, 288, 299, 300, 304, 319, 320, 321, 322, 323, 324 |
| Strep•ADP-Hep•FDLA•HepI | Arg60Ala | 1.93 ± 0.23 | 21.17 ± 0.16 | 29.93 ± 0.50 | 59, 60, 61, 62, 63, 64, 65, 66, 67, 68, 69, 70, 71, 72, 207, 317, 318, 319, 320, 321, 322, 323, 324 |
| Strep•ADP-Hep•FDLA•HepI | Arg61Ala | 1.88 ± 0.34 | 21.21 ± 0.14 | 30.10 ± 0.38 | 63, 64, 65, 66, 67, 68, 69, 70, 71, 72, 73, 75, 76, 79, 83, 157, 187, 189, 207, 280, 281, 288, 299, 300, 301, 317, 318, 319, 320, 321, 322, 323, 324 |
| Strep•ADP-Hep•FDLA•HepI | Arg63Ala | 1.76 ± 0.22 | 21.26 ± 0.16 | 30.15 ± 0.37 | 62, 63, 64, 65, 66, 67, 68, 69, 70, 71, 72, 73, 75, 83, 157, 207, 280, 281, 284, 288, 298, 299, 300, 301, 302, 303, 304, 305, 317, 318, 319, 320, 321, 322, 323, 324 |
| Strep•ADP-Hep•FDLA•HepI | Lys64Ala | 1.61 ± 0.19 | 21.12 ± 0.12 | 29.80 ± 0.36 | 63, 64, 65, 66, 67, 68, 69, 207, 231, 300, 319, 320, 321, 322, 323, 324 |

Supplemental Table 2: Average values of RMSD, and radius of gyration, interdomain center of mass distance. RMSF values are reported for residues that have a greater than 1.5 Å fluctuation.

| Inhibitor | Mutation | $\Delta G_{Exp}$ (kcal/mol) | $\Delta G_{Calc}$ (kcal/mol) | $\Delta\Delta G_{Exp}$ (kcal/mol) | $\Delta\Delta G_{Calc}$ (kcal/mol) | Pearson Correlation Coefficient (R) |
|---|---|---|---|---|---|---|
| Tobramycin | WT | -8.1 ± 0.1 | -10.5 ± 5.7 | --- | --- | 0.87 |
| | Arg60Ala | -7.5 ± 0.1 | -8.1 ± 2.3 | 0.7 ± 0.1 | 2.4 ± 6.2 | |
| | Arg120Ala | -8.7 ± 0.2 | -13.8 ± 3.5 | -0.6 ± 0.2 | -3.3 ± 6.7 | |
| Streptomycin | WT | -8.5 ± 0.1 | -13.1 ± 5.3 | --- | --- | |
| | Arg60Ala | -7.1 ± 0.4 | -9.9 ± 3.1 | 1.4 ± 0.4 | 3.2 ± 6.1 | |
| | Arg61Ala | -7.0 ± 0.2 | -10.9 ± 2.1 | 1.5 ± 0.2 | 2.2 ± 5.7 | |
| | Arg63Ala | -7.1 ± 0.1 | -8.5 ± 1.5 | 1.4 ± 0.1 | 4.6 ± 5.5 | |
| | Lys64Ala | -7.1 ± 0.1 | -6.8 ± 7.6 | 1.3 ± 0.1 | 6.3 ± 9.2 | |

Supplemental Table 3: (**A**)Table of binding energies of Streptomycin/Tobramycin to HepI both wildtype and Arg/Lys mutants with correlation values.

### End of Supplemental
